# Single-cell morphology encodes functional subtypes of senescence in aging human dermal fibroblasts

**DOI:** 10.1101/2024.05.10.593637

**Authors:** Pratik Kamat, Nico Macaluso, Chanhong Min, Yukang Li, Anshika Agrawal, Aaron Winston, Lauren Pan, Bartholomew Starich, Teasia Stewart, Pei-Hsun Wu, Jean Fan, Jeremy Walston, Jude M. Phillip

## Abstract

Cellular senescence is an established driver of aging, exhibiting context-dependent phenotypes across multiple biological length-scales. Despite its mechanistic importance, profiling senescence within cell populations is challenging. This is in part due to the limitations of current biomarkers to robustly identify senescent cells across biological settings, and the heterogeneous, non-binary phenotypes exhibited by senescent cells. Using a panel of primary dermal fibroblasts, we combined live single-cell imaging, machine learning, multiple senescence induction conditions, and multiple protein-based senescence biomarkers to show the emergence of functional subtypes of senescence. Leveraging single-cell morphologies, we defined eleven distinct morphology clusters, with the abundance of cells in each cluster being dependent on the mode of senescence induction, the time post-induction, and the age of the donor. Of these eleven clusters, we identified three *bona-fide* senescence subtypes (C7, C10, C11), with C10 showing the strongest age-dependence across a cohort of fifty aging individuals. To determine the functional significance of these senescence subtypes, we profiled their responses to senotherapies, specifically focusing on Dasatinib + Quercetin (D+Q). Results indicated subtype-dependent responses, with senescent cells in C7 being most responsive to D+Q. Altogether, we provide a robust single-cell framework to identify and classify functional senescence subtypes with applications for next-generation senotherapy screens, and the potential to explain heterogeneous senescence phenotypes across biological settings based on the presence and abundance of distinct senescence subtypes.

## INTRODUCTION

Senescence represents a heterogeneous cellular phenotype characterized by stable proliferative arrest^1,2^, upregulation of cyclin-dependent kinase (CDK) inhibitors such as p16 and p21^3^, increased secretions of pro-inflammatory molecules^4^, and drastic changes in cell and nuclear morphologies^5–7^. Although these senescence-associated changes have been critical to shaping our current understanding of senescence phenotypes across cell populations, it is limited. These limitations stem from the notion that: i) there are currently no absolute biomarkers of senescence, but as a field we use a handful of protein-based biomarkers and more recently curated gene expression signatures^8,9^ as proxies of senescence^1,3^, which in many cases do not capture all senescent cells across biological settings^10^. ii) Cellular senescence is typically defined as a binary phenotype, meaning that cells are considered as senescent of not. Furthermore, this binarization comes with the implied assumption that senescent cells are in some ways equivalent and may respond similarly to biological perturbations, such as senotherapies. However, mounting evidence is demonstrating that senescence is heterogeneous and context-dependent, conferring both favorable and deleterious effects based on the biological setting^3,11^. Addressing these limitations are key to developing a deeper understanding of cellular senescence and the factors that determine their heterogeneous phenotypes.

Significant advances in single-cell technologies and modern data science approaches have led to novel methods to profile senescent cells^5–7^. Some studies using next-generation sequencing technologies on bulk or single-cell populations have demonstrated robust ways to profile large numbers of heterogeneous senescent cells based on curated gene sets^8,9^. These approaches have enabled the simultaneous interrogation of multiple transcriptionally-defined markers and putative molecular signatures. Within human skin, some of these studies have identified multiple subpopulations of fibroblasts with differential gene expression patterns^12^, leading to questions of whether these subpopulations exhibit differential senescent phenotypes and responses to senescence induction, and their potential roles in pathogenesis and aging. Using a WI-38 fibroblast cell line, another study demonstrated the emergence of senescence clusters based on differential gene expression patterns of single cells, which was shown to depend critically on the mode of senescence induction^13^. While it is fascinating that transcriptionally-defined senescent subtypes exist and that they depend on how senescence was induced, performing single-cell sequencing brings a few challenges related to the high cost of sequencing and the throughput and feasibility of profiling multiple samples and conditions. Taken together, these advances point to the additional need to develop robust, cost-effective, and high-throughput approaches to profile senescence cells.

Recent image-based strategies have employed high-content microscopy coupled with machine learning to profile senescent cells cultured on flat substrates coated with physiologically-relevant extracellular matrix (ECM) proteins. These studies used microscopy images of senescent and non-senescent cells to train computational models to determine whether a cell was senescent or not, with downstream validation of senescence based on the expression of senescent biomarkers^5–7^. Furthermore, coupling high content imaging with machine learning is a powerful tool to identify senescence with potential applications to screen libraries of repurposed drugs for identifying new senotherapies or compounds capable of inducing senescence^6,7^. Additionally, one of the studies demonstrated that their model trained on nuclear features of cultured senescent cells can be directly applied to identify senescent cells within tissue sections^5^. While these recent studies provide a proof-of-concept for identifying senescence cells using imaging, these studies made fundamental assumptions related to i) senescence being a binary phenotype (*i.e.,* cells are senescent or not), and ii) that all cells exposed to senescence inducers were senescent and vehicle-treated cells were non-senescent^5–7^. While assumptions like these are potentially true in some biological contexts, generalizations under these assumptions could bias the interpretation of results. Taken together, image-based approaches coupled with machine learning provides a powerful solution to identifying senescent phenotypes, but there is a critical need to go beyond binary classifications to investigate induction kinetics and the presence of putative senescence subtypes.

Here we present an integrated image analysis and machine learning workflow for identifying and quantifying senescence, which we call SenSCOUT (***Sen****escent **S**ubtype **C**lassifier based on **O**bservable **U**nique pheno**t**ypes*), to identify and classify functional subtypes of senescence, with specific emphasis on mitigating some of the current limitations with senescence identification and classification detailed above. Using primary dermal fibroblasts from two donors aged 23 and 89 we initially profiled the morphologies of cells at baseline, after serial passaging (replicative senescence) and eight days post-induction using four chemical inducers. By profiling single cells across each of these biological conditions, we identified eleven morphologically-defined cellular subtypes out of which we denoted three as *bona fide* senescence subtypes (C7, C10, C11) based on the exhibited morphologies and the expression of multiple protein-based senescence biomarkers (up to five biomarkers). As part of this workflow, we also implemented and validated a label propagation scheme using imputation based on a k-nearest neighbor (knn) approach. This imputation approach enabled us to quantify and validate the expressions of protein-based biomarkers per cell based on their morphologies.

In addition to the parameterization of the cellular and nuclear morphological features, we developed a machine learning scheme based on the Xception^14^ architecture to classify cells and quantify a senescence score per cell based on raw textured images of F-actin (cell) and DNA (nuclei). Using this senescence score, we showed that the senescence burden across cell populations were dependent on the mode of senescence induction, time post-induction, and the age of the donor. To establish the association between senescence, senescence subtypes, and age, we used primary dermal fibroblasts from an expanded cohort of fifty healthy donors ranging in age from 20-89 years old. We observed that not only did the amount of senescent cells increase with age, but that senescence subtype C10 showed the strongest age-dependence. Furthermore, we observed that the baseline senescence burden was indicative of the senescence score after exposure to senescence-inducer doxorubicin. Lastly, to determine whether the identified senescence subtypes were functionally different we profiled the responses of senescence cells to a small panel of senotherapies with focused analysis on responses to Dasatinib + Quercetin (D+Q). Results showed that senescent cells responded based on four dominant patterns: short death, long death, stable viable, and unstable viable. Senescent cells treated with D+Q exhibited higher abundance in short death, long death, and unstable viable, with cells in C7 being the most sensitive to death after D+Q exposure.

Altogether, we provide a robust single-cell framework using imaging, morphological profiling, and machine learning to identify and classify functional subtypes of senescence across multiple biological conditions.

## RESULTS

### Comprehensive profiling of cellular senescence across multiple induction conditions and biomarkers

Cellular senescence can be induced across *ex vivo* cell populations based on a variety of methods that take advantage of core mechanisms such as DNA damage responses (DDR), replication stress and telomere shortening, oncogenic activation (*i.e.,* p53 and RAS), and epigenetic modifications^15^. To recapitulate the range of physiologically-relevant senescence phenotypes, we established a framework using primary dermal fibroblasts isolated from a 23- and an 89-year-old, referred to as young and old, respectively. To profile senescence, we exposed cells from both donors to optimized concentrations of four chemical inducers (*i.e.,* Bleomycin (BLEO), Doxorubicin (DOX), Atazanavir (ATV), Hydrogen peroxide (H_2_0_2_)), and serially passaged cells (REP) until proliferation arrest (see Methods). After induction, the senescence phenotype was allowed to develop over the course of eight days, after which cells were plated at low density onto Collagen-I coated glass bottom dishes, fixed and stained for proteins/biomarkers of interest, then imaged for downstream single-cell morphology analysis (**Figure 1a**). For all experiments reported in this study, unless otherwise stated, cells were stained with Hoechst 33343 (H33342) to delineate nuclear boundaries, Phalloidin to delineate cell boundaries, senescence biomarker p21, and either Lamin B1 (LMNB1), High Mobility Group Box 1 (HMGB1), p16, or beta-galactosidase (**Figure 1b, Supplementary Figure 1a**) as additional biomarkers of senescence^3,16^. To quantify single-cell morphologies, baseline uninduced cells from both donors and the senescence-induced cells were segmented using an optimized CellProfiler pipeline^17^ then we implemented an in-house post-processing pipeline for data curation (*i.e.,* removing mis-segmented cells and outliers) and analysis (see Methods).

**Figure 1.**
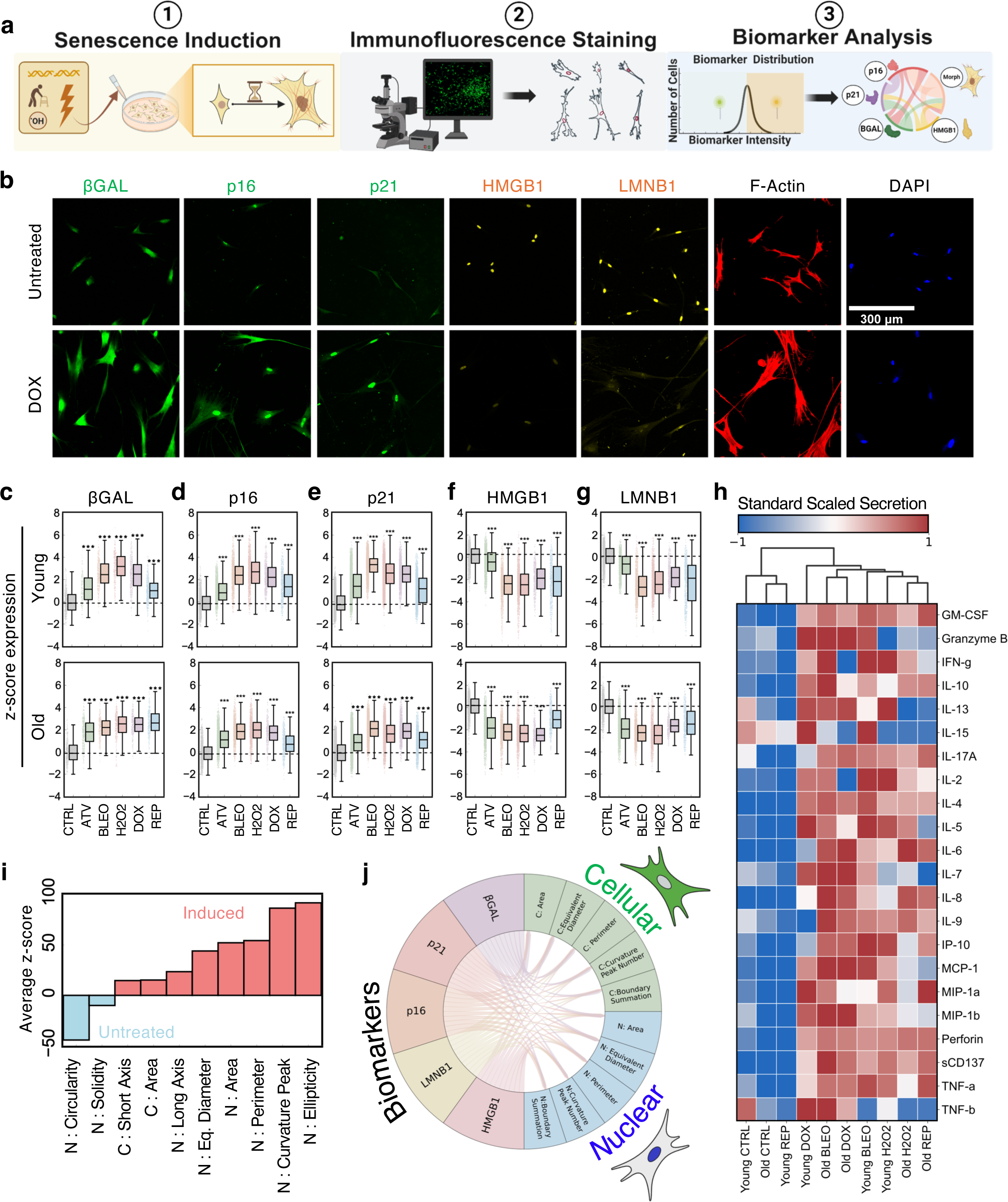
Cellular and nuclear morphology are biomarkers for in vitro cellular senescence. **a.** Graphical illustration of experimental workflow to induce and characterize senescent cells. **b.** Representative images of the immunofluorescent staining of 5 senescence-associated protein biomarkers across untreated (DMSO vehicle) and DOX-induced primary dermal fibroblast samples (age 23-male). **c-g.** Z-score fluorescence quantification relative to control of βGal (c), p16 (d), p21 (e), HMGB1 (f), and LMNB1 (g) for young and old dermal fibroblast samples (*n*>400 single cells per condition, mean ± 95% CI, One-Way ANOVA relative to age-matched control expression, p ≤ 0.001 relative to control). **h**. Heatmap of standard scaled expression of pro-inflammatory secretions for induction conditions (average algorithm, Euclidean linkage). **i**. Waterfall plot of cellular and nuclear morphological features enrichment to either untreated or senescence-induced samples across both ages (cutoffs= Log_2_ > 10). **j**. Circos plot of the 5 stained senescence-associated protein biomarkers and highly correlated morphological features. Connecting arcs represent an absolute value correlation coefficient greater than 0.3.

Altogether, we generated an initial dataset consisting of >50,000 single-cells, each having information on cell and nuclear morphologies with the expression of multiple protein-based senescence biomarkers for both ages across five senescence-induction conditions.

### Establishing single-cell senescence phenotypes across multiple biological conditions

To provide in-depth profiling of cells across baseline uninduced and senescence-inducing conditions, we quantified the expression of co-stained senescence biomarkers, cellular secretions, and parameters describing the average morphologies for populations of cells and their corresponding nuclei. To establish and validate the senescence phenotypes we first evaluated whether there was a halt in proliferation for senescence induced cells relative to the baseline uninduced cells (**Supplementary Figure 1a-b**). Given the observed proliferation arrest of senescence induced cell populations, we then quantified the cellular expressions based of protein-based senescence biomarkers in cells from both donor across five biological conditions. As expected, we observed an increase in the average expression of senescence biomarkers beta-galactosidase, p16, and p21, and a decrease in the expression of HMGB1 and LMNB1 for both donors relative to their baseline uninduced cells (**Figure 1c-g**). Interestingly, we observed that the extent of increase or decrease in the biomarker expressions was dependent on the mode of senescence induction and the donor age. Together, these results suggest that the mode of senescence induction matters, and that it contributes to differential abundances of senescent and non-senescent cells per condition.

To further establish the senescence phenotype, we profiled the cellular secretions at baseline and eight days post induction. We performed secretion analysis using the Bruker CodePlex^TM^ multi-analyte secretory panel, which contained overlapping analytes (cytokines, chemokines, etc.) to those identified in the core senescence associated secretory phenotype (SASP^4,18^), such as IL-6, IL-7, IL-8, IL-15, and MIP-1a. The abundance of secreted factors was quantified from the conditioned media across each condition by culturing a known number of cells for 48 hours, harvesting the conditioned media, and profiling the un-diluted conditioned media. From this analysis, we profiled twenty-two secreted factors across ten condition. Results showed that uninduced baseline controls for both ages clustered together and were defined by low expression of SASP-related and other pro-inflammatory proteins (**Figure 1h**). The senescence induced conditions on the other hand, clustered together with elevated expressions of SASP-related proteins. Consistent with the protein-based biomarkers, we also observed induction-dependent trends in the secretions. For instance, IL-7 was highly secreted by cells from both the young and old donor treated with DOX but was expressed at a lower level when cells were exposed to H2O2. Together, analysis of cellular secretions provides additional confirmation of bulk senescence phenotypes across our tested induction conditions.

Lastly, given the fact that we are collecting microscopy images of cells and nuclei across all conditions, we quantified the senescence-associated changes in cell and nuclear morphologies. Consistent with the published literature, we observed a significant increase in the sizes of both cells and nuclei post-induction (**Figure 1i**). For each cell within our dataset, we quantified over 250 morphological parameters, which we reduced using factor analysis to 90 morphological parameters that described the size, shape, curvature, and roughness of both cells and nuclei (see **Methods, Supplementary Table 1**). To push our analysis beyond the increase in cell and nuclear size, we asked which morphological features were most strongly associated with uninduced cells and senescence-induced cells, respectively. Performing differential-morphology analysis on the induced and uninduced cells, we observed that the majority of the highly differential morphological features, on both sides, were associated with nuclear morphologies (**Figure 1i**). However, features describing nuclear shapes were more strongly associate with the uninduced cells and features describing the nuclear sizes were more strongly associated with senescence-induced cells (**Supplementary Figure 1c**). These results fit well with recent findings from others showing that nuclear features strongly encode senescence information^5,7^. Given this associate between cell and nuclear morphology, biomarker expressions, and senescence, we wondered whether cell and nuclear morphologies were correlated with expression features of senescence biomarkers. To assess this, we performed correlation analysis across all experimental conditions for the 90 cell and nuclear parameters and the biomarker expressions. Interestingly, we observed multiple correlations among features given a threshold of 0.3 for the Pearson correlation coefficient (**Figure 1j**).

Taken together, we show using multiple lines of evidence that the biological conditions used in this study yields consistent senescence phenotypes with current literature, thereby providing the basis to develop our single-cell framework of senescence and test the notion of functional senescence subtypes.

### Single-cell morphologies encode heterogeneous senescence phenotypes in primary dermal fibroblasts

Given the capacity of cell and nuclear morphologies to encode senescence phenotypes, we developed a single-cell framework to identify and classify morphological subtypes of senescent and non-senescent dermal fibroblasts. To establish this single-cell framework cells from both donors across uninduced and senescence-induced conditions were imaged using high content microscopy, then cell and nuclear boundaries were segmented using an optimized pipeline in CellProfiler. Once segmented, we computed parameters that described cell and nuclear morphologies, as well as the expression of two protein-based senescence biomarkers (p21 and a combination of either p16, beta-galactosidase, HMGB1, or LMNB1; see Methods).

With this data, we pooled cells from twelve biological conditions, consisting of cells from the young and older donor at baseline (uninduced) or post-induction with optimized concentrations and durations of ATV, DOX, BLEO, H2O2, and REP (**Figure 2a**). Data visualization using two-dimensional UMAP revealed specific cell localization patterns per condition, with cells in uninduced conditions localizing to the lower left of the manifold and cells from induced conditions localizing towards the right of the manifold (**Figure 2b**). Results also showed a higher spread of the uninduced cells from the old donor towards the right of the manifold, suggesting a slightly higher abundance of senescent-like cells relative to the young donor. Interestingly, although different inducers exhibited varying degrees of spread across the manifold, there were core overlap areas for all inducers. To gain a deeper understanding of these spatial patterns across the manifold, we performed k-means clustering analysis to identify morphologically distinct groups of cells. K-means analysis revealed eleven distinct morphological clusters, each having characteristic cell and nuclear morphologies (**Figure 2c**, **Supplementary Figure 2b-c**). Importantly, visual inspection of the cellular morphologies indicated a pattern of increasing size from left to right and increasing linearity of the shapes from top to bottom of the manifold. These results were confirmed by color-coding the magnitude of key morphological parameters onto the UMAP, with increased cell and nuclear area from left to right and increasing cell aspect ratio (corresponding decrease in cell circularity) from top to bottom (**Figure 2d**). Of note, cluster C9 contained a large fraction of mis-segmented cells, as such we kept C9 as part of our quality control step (mis-segmented cluster) but omitted cells classified in C9 from formal analysis and reporting in the results.

**Figure 2.**
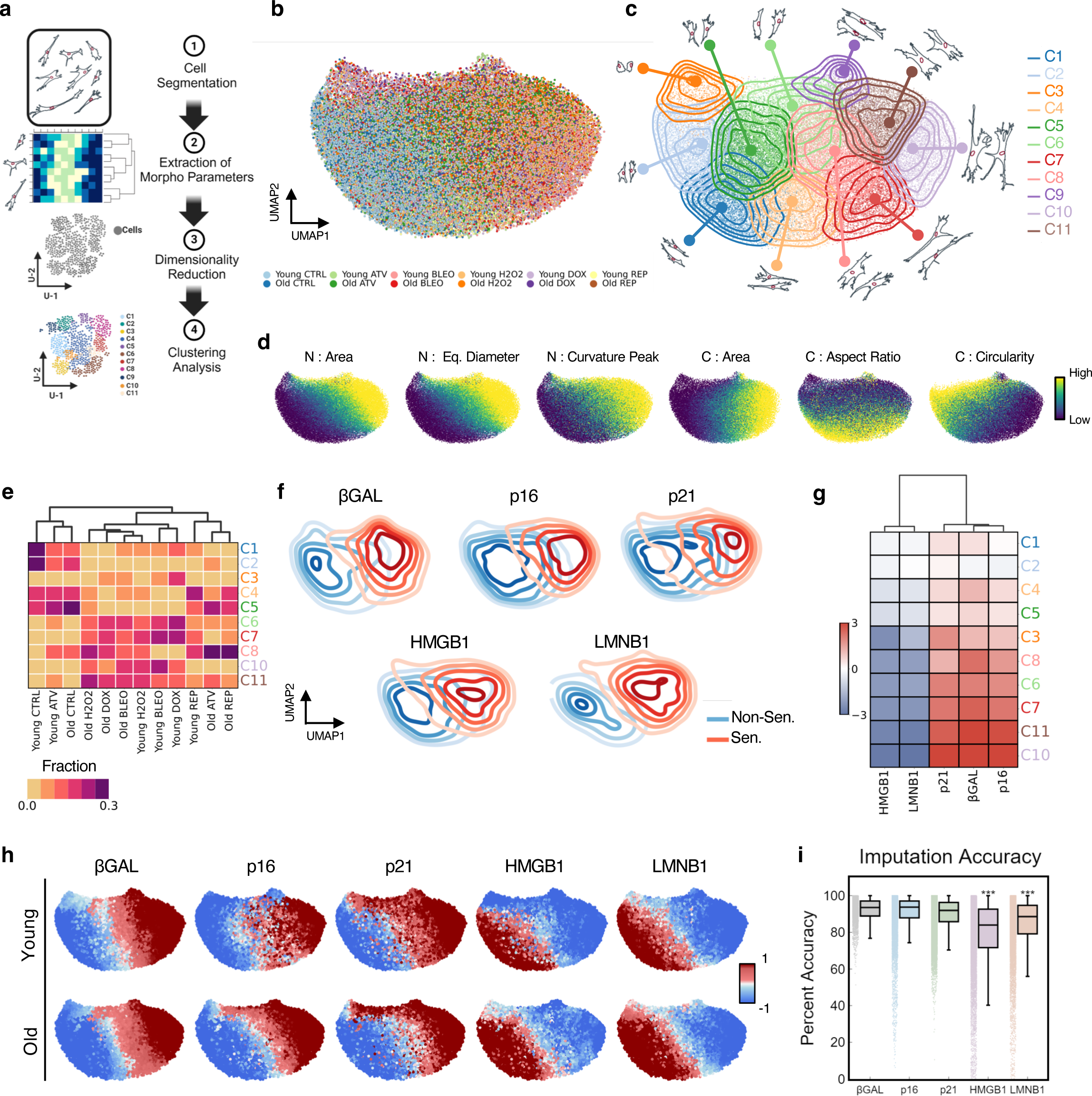
Morphology encodes heterogeneity within the senescent phenotype. **a.** Schematic of morphological analysis pipeline. **b.** Scatter of UMAP space based on 87 morphological parameters for all conditions across both age groups (n≈48,000 single cells). Individual points represent single cells. **c.** Overlay of the 11 optimal KMEANS clusters on top of the UMAP space with representative cellular (outer) and nuclear (inner) morphologies. Contour lines indicate gaussian kernel density function for each cluster. Cluster 9 consisted of mis-segmented cells and was dropped from subsequent analysis **d.** Plots of select morphological features layered on top of the UMAP space; navy and yellow colors delineate high and low standard scaled expression, respectively. Generally, UMAP-1 (x-axis) was correlated with nuclear and cellular size and UMAP-2 (y-axis) was correlated with cell rounding. **e.** Heatmap displaying the fractional abundance of cells within each KMEANS cluster for each experimental group (*n*>1000 cells per condition, average algorithm, Euclidean linkage). **f.** Contour overlay of hypothesized non-senescent and senescent classifications for βGal, p16, p21, HMGB1, and LMNB1. Senescence cutoff was determined by the z-score magnitude being greater than 2 with respect to upregulation or downregulation of the biomarker. **g.** Heatmap of the average quantified protein biomarker expression per cluster, organized top-bottom to match left-right traversal across the manifold (z-score expression, average clustering algorithm, Euclidean linkage). **h.** Scatter plots of the UMAP space with biomarker expressions across young and old samples. Visualization uses a combination of ground truth (if applicable) and imputed biomarker expressions to fill in the breadth of the manifold. **i.** Imputation error using morphological parameters of single cells to predict protein biomarker expression (n>500 cells per biomarker, mean ± 95% CI, multiple comparison Tukey test, p ≤ 0.001 for HMGB1 and LMNB1 relative to other biomarkers). Accuracy is defined as the one minus the percent deviation from model-predicted value divided by the ground truth value. HMGB1 and LMNB1 expressed error statistically difference compared to all other biomarkers.

With these morphological clusters, we then assessed the presence and abundance of cells per morphological clusters across induction conditions (**Supplementary Figure 2d-e**). Unsupervised hierarchical clustering of induction conditions per cluster revealed the emergence of three groups based on the dendrogram branches. With uninduced conditions for young and old donors and young cells induced with ATV consisting group 1, both young and old cells induced with DOX, BLEO, and H2O2 consisting another group 2, and young and old cells induced via serial passaging (REP) and old cells induced with ATV consisting the last group 3 (**Figure 2e**). Interestingly, group 2 containing young and old cells treated with DOX, BLEO, and H2O2 exhibited enrichments in clusters C6, C7, C8, C10, C11, whereas group 1 were depleted in those clusters but enriched for C1, C2, C4, and C5. Group 3 on the other hand, contained a mixture of enriched morphological clusters, with noted overlaps with both group1 and group 2, and general enrichments in C4, C5, and C8. Furthermore, we noticed that induction conditions exhibited higher heterogeneity based on the Shannon entropy relative to uninduced conditions for both ages, suggesting that induced cells exhibited a wider range of morphologies relative to uninduced cells (**Supplementary Figure 2f**).

Taken together, we present a robust framework to define senescence phenotypes across multiple biological conditions. Given these morphology-defined clusters, we can now test whether these morphological clusters reveal functional subtypes of senescent and non-senescent cells.

### Profiling single-cell morphologies based on senescence biomarkers

Current approaches to identify senescence across cell populations rely on profiling the presence and expression of protein-based biomarkers, including p16, p21, HMGB1, LMNB1, beta-galactosidase^19–21^. For instance, senescence cells tend to express increased expression of p16, p21, beta-galactosidase, and decreased expression of HMGB1 (translocation out of the nuclei) and LMNB1 relative to non-senescent cells. While these expression patterns have been collectively established across multiple studies, many single studies evaluating senescence tend to profile one or maybe two of these biomarkers. To provide a comprehensive assessment of the expression patterns across the biological conditions tested, we profiled all five of the biomarkers listed above. Since our imaging workflow was only capable of imaging four fluorescence channels at a time, we devised a multiplexed strategy to stain p21 plus one of the remaining four biomarkers. This was iterated across all samples, and biological replicates were stained with different combinations until we had all five biomarkers represented across each of the twelve biological conditions (see Methods). With this workflow, we then quantified the localization of cells expressing elevated or depleted levels of the senescence biomarkers within our UMAP space.

To determine the thresholds per biomarker for positive and negative expression (*i.e.,* positive = senescent), we first plotted and overlaid the distributions of single-cell expressions in the uninduced and senescence-induced conditions. Then we identified the thresholds on the basis that senescence cells will be enriched in the induced conditions and have a low representation in the uninduced cell population. Practivally speaking, we defined thresholds per biomarker based on the magnitudes of the shifts in distribution post induction (see Methods). Once we determined these thresholds, we next investigated how cells expression high and low levels of each protein-based biomarker of senescence localized across the manifold and the morphological profiles across each cluster. In essence, we quantified the presence and spread of “hot” and “cold” zones, denoting senescent and non-senescent cells. Results showed a clear separation of cells classified as senescent (red contours) and non-senescent (blue), with senescence emerging from left to right of the manifold (large cells with large nuclei were localized to the right), confirming qualitative observations presented earlier (**Figure 2f**). Interestingly, while the contours showed clear overlap areas among all senescence biomarkers, the unique topologies of each biomarker suggest that some biomarkers may identify cells as senescence that others may missed. For instance, the contours for senescent cells identified by p16, beta-galactosidase, and LMNB1 had less spread across the manifold relative to p21 and HMGB1, meaning that HMGB1 and p16 will identify more cells as senescent relative to using p16, LMNB1 or beta-galactosidase. Whether p21 or HMGB1 is more accurate remains to be determined, however, the union of the biomarkers likely best captures the senescence phenotypes.

Next, we quantified the average expression of each senescence biomarker per cluster. Results showed that clusters C7, C10, and C11 had high relative expressions of p21, p16, and beta-galactosidase and low expression of HMGB1 and LMNB1, suggesting an enrichment for senescence within those clusters. Clusters C1, C2, C4, and C5 had the opposite expression pattern, suggesting that they were enriched for non-senescent cells. Lastly clusters C3, C6, and C8 had intermediate expressions of the senescence biomarkers, which suggests that those clusters harbored an intermediate state or cells that may be potentially ‘poised’ toward senescence (**Figure 2g**). In summary, these findings highlight a strong relationship between cell and nuclear morphologies and the expression of senescence biomarkers across our morphology-defined clusters, underscoring the notion that morphology robustly encodes the heterogeneous senescence phenotypes.

### Single-cell morphology robustly encodes senescence biomarker expressions

Given this robust link between single-cell morphologies and senescence biomarker expressions, we wondered whether we could use cellular morphologies to directly predict the biomarker expression for all five biomarkers per cell. To test this, we developed and implemented an imputation-based label propagation scheme using k-nearest neighbors^22,23^. Briefly, we constructed the imputation scheme by leveraging cell and nuclear morphologies to determine the senescence biomarker expressions based on the proximity to cells having known expressions of the biomarker of interest (**Supplementary Figure 2g**). For instance, to impute the expression of p16 in a cell stained for p21 and LMNB1 (not stained for p16), we identified the 20 nearest cellular neighbors that had measured expression of p16, then weighted the expressions based on the Euclidean distance from each of the 20 neighbors (see Methods). This was iterated for every cell across all senescence biomarkers until we had an expression profile for each of the five senescence biomarkers per cell. Meaning that each of the 50,000 single cells had two measured and three imputed biomarker expression profiles (**Figure 2h**). Results from the imputation scheme proved to be robust, with prediction accuracies over 85% for each of the senescence biomarkers relative to ground truth expressions (*i.e.,* the nearest cells with measured expression of the biomarkers) (**Figure 2i**).

Somewhat unexpectedly, we observed that some cells had lower imputation accuracies, particularly for HMGB1 and LMNB1; beta-galactosidase, p21 and p16 had fewer of those cells. We rationalized that this may be attributed in part by those biomarkers having a lower dynamic range for senescent versus non-senescent cells, and that the distances of cells from a neighbor with known expression was larger. However, even with the subset of cells having lower accuracies, the pattern of localization for the imputed cells across the manifold was consistent across both ages and for cells with measured expression per senescence biomarker (**Figure 2h**).

Given this comprehensive dataset with all five senescence biomarkers per cell, we considered how the biomarkers interfaced with each other and whether certain biomarkers identified senescence better than others. To answer this, we took all cells, across all biological conditions and grouped cells based on the number of senescence biomarkers that were positive based on the previously described thresholding strategy. Then within each of the comparisons (*i.e.,* 2, 3, 4, or 5 senescence biomarkers) we quantified the fraction of cells with each of the biomarkers (**Supplementary Figure 2h**). We found that for all the cells having two positive senescence biomarkers, approximately 50% of cells were classified as senescent based on p21 and HMGB1. Interestingly, of the cells classified as senescent based on three senescence biomarkers, over 70% were positive to beta-galactosidase, HMGB1 and LMNB1. Furthermore, of the cells considered senescent based on four biomarkers, beta-galactosidase, HMGB1, and LMNB1 were expression in more than 80% of them, with p16 being the forth biomarker having expression in approximately 70% of the cells. Within this framework, we also mapped the localization of cells with varying number of senescence biomarkers from zero (*i.e.,* non-senescent cells) to five (senescence based on all five biomarkers). Qualitative results recapitulated the shift from left to right from zero to five markers, with a tight spread corresponding to the overlap areas across all the biomarkers (**Supplementary Figure 2i**).

Taken together, we show that information on the protein-based senescence biomarker expressions are nested within the senescence-associated morphologies, with the capacity to impute the biomarker expressions based on cell and nuclear morphologies.

### Machine learning strategies reveal morphological subtypes of senescence

While we were able to establish robust associations between cell and nuclear morphologies and senescence phenotypes at single-cell resolution, these findings were generated from parameterizing cell and nuclear morphologies based on size, shape, curvature, and boundary roughness (**Supplementary Table 1**). To extract information that may not be captured by parametrized morphological features, we implemented a machine learning scheme using the raw textured images of cells (stained F-actin) and nuclei (stained DNA) across individual cells, with the goal of generating a senescence score per single cell. First, we constructed a three-layer data instance for each individual cell, where the first layer was the cell’s nuclei, the second layer was the cell’s F-actin structure, and the final layer consisting of a composite image both the F-actin and nuclei structures. Each of these data instances per cell were cast onto a blank canvas having dimensions of 1526 by 1526-pixel, with the geometric centroid of each object (nuclei, cell, composite) mapping to the center of the canvas. We hypothesized that each data instance captured not only the ensemble of possible morphologies associating with senescence, but that it also offered unbiased insights into which properties were most important to the senescence identification and classification. Each data instances were labeled as either ‘non-senescent’ or ‘senescent’ based on the thresholded expressions of at least two senescence biomarkers. We trained our model using the Xception^14^ architecture with a SoftMax output^5^ (**Figure 3a**). The SoftMax output, ranged from ‘0’ to ‘1’ based on the probability of a cell being non-senescent or senescent according to the predefined threshold biomarker expressions. We implemented a balanced cohort of 6000 data instances spanning all induction conditions, both ages, and all five senescence biomarkers, with the model trained for 220 epochs. The trained model had an average accuracy of 89% for both the training, test, and validation datasets **(Figure 3b, Supplementary Figure 3a-b)**. To further demonstrate the robustness of the model, we plotted the Receiver Operator Curve (ROC), which yielded an area under the curve (AUC) of 0.95 (**Figure 3c**).

**Figure 3.**
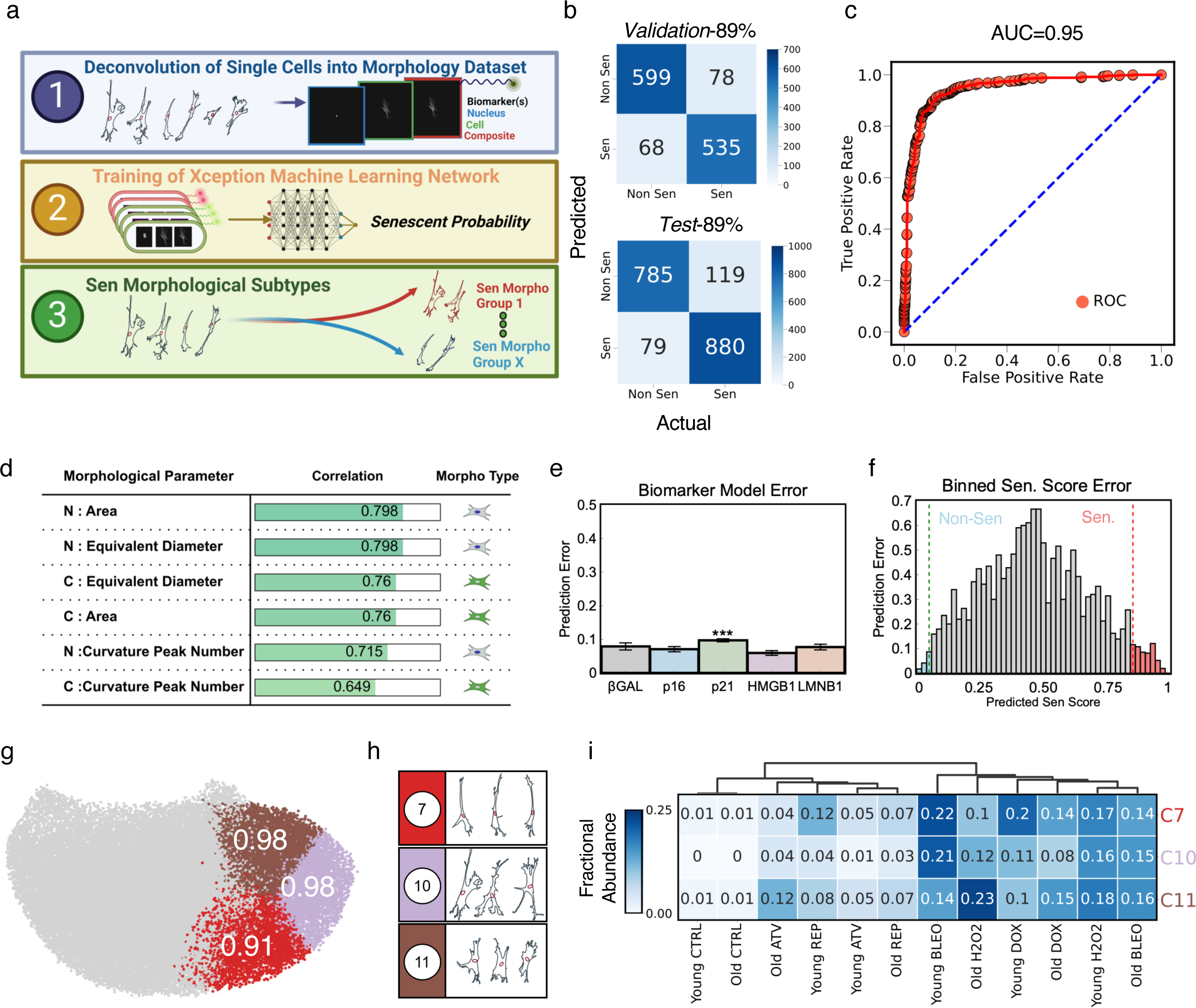
Machine learning and computational techniques identify morphological subtypes of senescence. **a**. Workflow for using cellular and nuclear textured images to develop single-cell resolution ‘senescent scores’, ranging from 0-1. Training was conducted on an age, induction, and biomarker balanced cohort of 6,000 single cells (see Methods for further detail on senescent cutoffs and dataset creation). **b**. Confusion matrix describing accuracy of trained Xception model on test and validation datasets. **c**. Receiver-operating characteristic (ROC) curve of the Xception model with an area under the curve (AUC) of 0.95. **d**. Top magnitude correlations between morphological parameters of single cells and their corresponding model-predicted senescent scores. **e**. Average model prediction error associated with each senescent-associated protein biomarker. Error is defined as the magnitude difference between predicted score and ground truth score (*n*>500 cells per biomarker, mean ± 95% CI, Tukey multiple comparison test, p ≤ 0.001 p21 relative to other biomarkers). **f**. Misclassification error associated with binned, model-predicted senescent scores (0.02 interval bins spanning 0-1). Error is defined as the fraction of cells within the bin in which the integer-rounded score does not match the true score (See Methods for further detail); bins with errors below 12% are highlighted in blue or red to highlight control and senescent high confidence regimes. **g**. Identification of 3 KMEANS morphological clusters (morphological subtypes) with average senescent scores within 12 % error. **h**. Representative morphologies of the three senescent morphological subtypes. **i**. Heatmap describing the differential enrichment amongst the three morphological senescence subtypes across all experimental conditions (average algorithm, Euclidean linkage).

In efforts to gain a deeper understanding of what morphological features the model was using to define senescence scores (probability of a cell being senescent based on two senescence biomarkers), we constructed a correlation matrix of the senescent score per cell versus the quantified morphological parameters used previously (**Figure 2**). Interestingly, we found strong correlations between senescence scores and parameters describing cell and nuclear sizes, with the Pearson correlation coefficients for nuclear area and equivalent diameter being approximately 0.8, and the same parameters of the cells being approximately 0.76. While these types of correlation analysis offer some insights to connect interpretable morphologies to the Xception model, the actual morphological features most weighted by the model may be poorly translated to the parametrized morphologies.

Previous studies using morphology to identify senescent versus non-senescent cells have often used either nuclear^5,7^ or cell^6^ morphology. While both have proven effective in capturing the senescence phenotype, there has been limited investigation using both cell and nuclear morphologies. To address this in an unbiased manner, we converted both training and test sets to include either nuclei or cell only. The resulting datasets were used to iteratively train new models for 50 epochs and we computed the accuracies based on protein-based biomarker expressions. Notably, both models resulted in statistically similar accuracies, indicating that both cell and nuclear morphologies both provide useful information on senescence phenotypes **(Supplementary Figure 3c)**. To further illustrate this, we computed the classification error between the senescence scores per cell and the senescence classification per biomarker. Results showed an error of less than 0.1 across all senescence biomarkers indicating that the Xception model accurately captured the senescence phenotype in an unbiased manner (**Figure 3e**).

While our trained model robustly quantifies senescence scores ranging from 0 to 1 for each cell, we wanted to determine a rigid threshold above or below which we could confidently determine a cell as senescent or not. Conventionally, binary SoftMax outputs are rounded to the nearest integer value for classification, but we wanted to quantify how specific scores correlate with the accuracy of the classification. In other words, how does the error of the classification change for a cell having a senescent score of 0.6 versus 0.7? To probe this, we binned the range of senescent scores between 0 and 1 with increments of 0.02 then assessed how many cells in each bin were correctly classified based on the biomarker expressions. From this analysis, we selected a score above 0.88 as senescent and below 0.1, corresponding to a prediction error below 12% (**Figure 3f, Supplementary Figure 3d**). To visualize how the model to determine senescence scores related to the morphology-defined clusters we projected the senescence score per cell across all clusters. Consistent with senescence phenotypes based on biomarker expressions per cluster (**Figure 2g**), we found that clusters C1, C2, C4, C5 exhibited low senescence scores (<0.3), C7, C10, C11 exhibited high senescence scores (>0.9), and C3, C6, C7, C8 exhibited intermediate scores (0.6-0.9) (**Supplementary Figure 3e**). Here we show that combining our Xception trained model with morphological parameters and senescence biomarkers provides a robust classification of senescence, with senescence scores increasing from left to right of the manifold (**Supplementary Figure 3f**) and clusters C7, C10, C11 exhibiting scores above 0.9 (**Figure 3g**) and large cell and nuclear morphologies (**Figure 3h**).

Lastly, given the high senescence scores for cells in C7, C10, and C11, we re-evaluated the idea that different induction conditions result in differential senescence phenotypes. Results indicated induction-dependent senescence phenotypes with varying abundance of cells in each of the three senescence clusters (**Figure 3i**). For instance, cells from the young donor induced with BLEO harbored 22% and 21% of cells in C7 and C10, respectively. Whereas cells from the old donor induced with H2O2 harbored 23% of cells in C11. Taken together, we show the robust identification and classification of senescence phenotypes bases on morphology, with validation metrics based on multiple protein-based senescence biomarkers. Furthermore, the identified morphology clusters exhibit varying senescence scores, with C7, C10 and C11 having a robust senescence phenotype. From now on we will refer to clusters C7, C10, and C11 as senescence subtypes.

### Profiling induction-specific kinetics to developing senescence phenotypes

While our previous analysis provides a robust framework to describe morphological patterns of cellular senescence, it was developed on snapshots at a single timepoint (day 8). To establish robust temporal patterns across senescence progression we performed senescence induction on cells from the young donor using DOX and H2O2, with snapshots of morphologies collected at multiple time points, specifically at 0, 2, 5, 8, 12, and 15 days post-induction. At each of these time points, we repeated the experimental and imaging workflows described previously (see Methods) to collect information on cell and nuclear morphologies, as well as the single-cell expressions of the five senescence biomarkers (**Figure 4a**). From this analysis, we observed differences in the senescence progression based on whether cells were treated with DOX or H2O2 (**Figure 4b-c**). For the DOX-induced condition, cells exhibited a steady progression towards a senescence phenotype, with a decrease of cells in C1, C2, C4, and C5 as a function of time post induction, and a complimentary increase of cells in C3, C6, C7, C10, and C11 up until day 15 (**Figure 4d, Supplementary Figure 4a, 4c**). H2O2-induced cells on the other hand exhibited a bi-phasic progression, with cells in C1, C2, C4, and C5 decreasing up to day 5 then steadily increasing again. Simultaneously, cells in C6, C7, C10 and C11followed the opposite pattern, with an increase up to day 5 and a decrease thereafter until day 15 (**Figure 4e**, **Supplementary Figure 4b, 4d**). Here the depletion of cells within clusters is not due to the cells dying, but due to a fractional redistribution of cells within clusters as they transform towards senescence. Furthermore, the apparent reversal of the senescence phenotype over time for cells treated with H2O2 is not that cells are going from senescent to non-senescent, but there is a small fraction of cells that do not transform and over time proliferates to recover the population.

**Figure 4.**
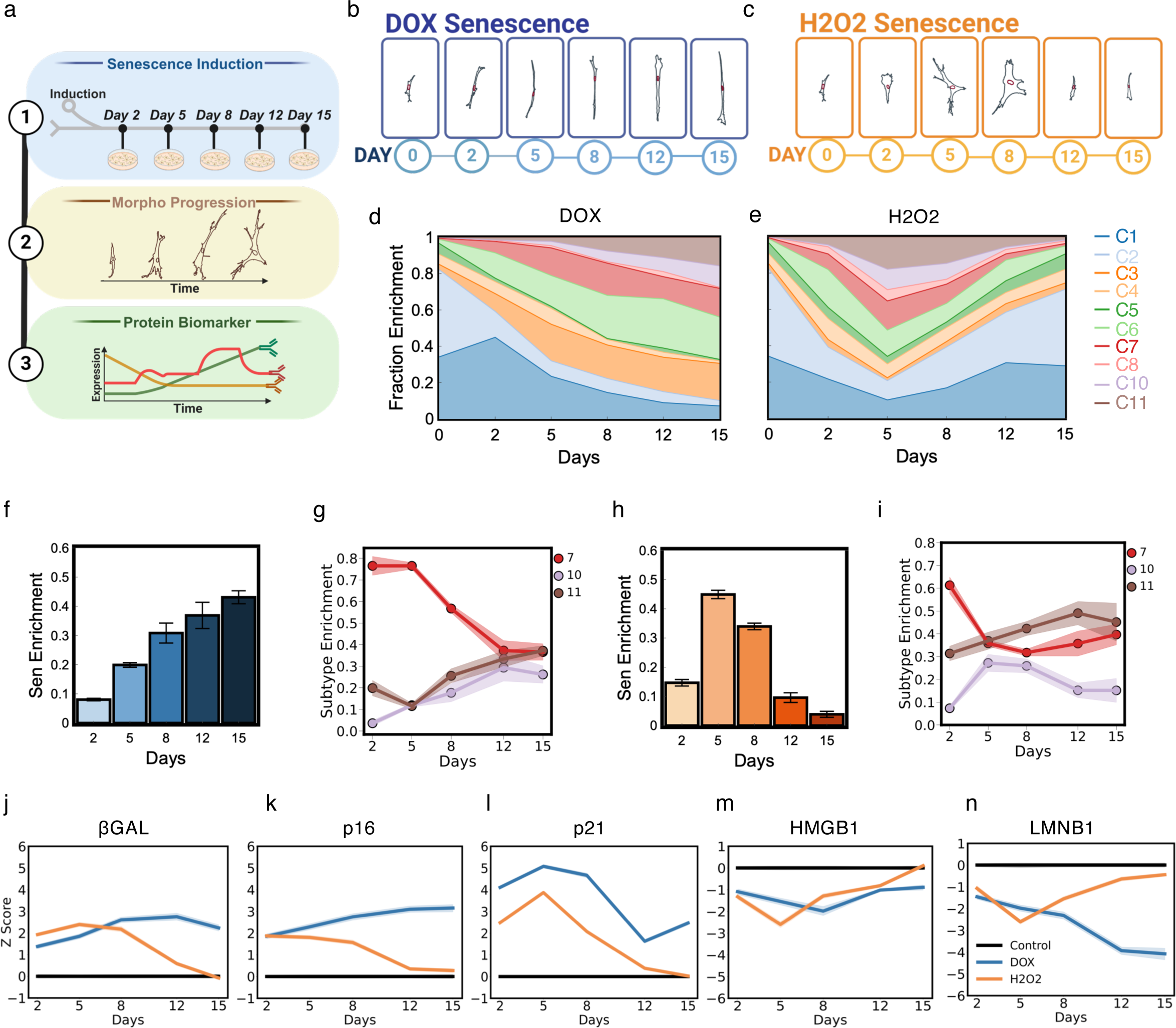
Senescence is a dynamic phenotype. **a.** Experimental design for analyzing the kinetics of senescence progression. **b-c.** representative morphologies for each time point for both DOX induction (b) and H2O2 induction (c). **d-g.** Senescence fraction enrichment and subtype dynamics with time. Box plots showing the fraction of cells within senescent morphological subtypes compared to all clusters for DOX (d) and H2O2 (f) induced samples (*n*>1000 cells per day, mean ± SEM). Line plots describe fractional distribution progression between the senescent morphological subtypes for DOX (e) and H2O2 (f) (shading indicates 95% CI around the mean). **j-n.** Average Z-score quantification of βGal (j), p16 (k), p21 (l), HMGB1 (m), and LMNB1 (n) as a function of time post senescence induction (shading indicates 95% CI around the mean). 23-year old fibroblasts were used for this analysis.

To quantify the trends in the senescence subtypes, we first quantified the senescence enrichments across the cell populations in both DOX (**Figure 4f**) and H2O2 (**Figure 4h**) induced cells. Confirming that in DOX, there was a steady increase in the senescence enrichment and a bi-phasic pattern for H2O2. Next, quantifying the temporal patterns for enrichment in each of the three senescence subtypes, we observed that for DOX there was a decrease in C7 and increases in C10 and C11 to convergence of approximately 30% in each subtype (**Figure 4g**), which was not as cleanly observed for H2O2 (**Figure 4i**). To help make sense of these results, we also quantified the temporal patterns of expression in each of the senescence biomarkers for both DOX and H2O2 conditions. Consistently, the results show that at the population level for each inducer that at early time point there is a deviation in the expression from the baseline uninduced cells. However, at later time points, H2O2 regains uninduced levels (**Figure 4j-n**). Taken together, results suggest that the temporal patterns of senescence are inducer-dependent, with cell populations fractionally redistributing among senescence subtypes during the progression. Furthermore, H2O2 may be a weaker inducer relative to DOX, with a fraction of the population not entering senescence and as a result the population recovers as the cells begin to proliferate at late time points.

### Profiling senescence subtypes and susceptibility as a function of chronological age

Multiple studies have reported the increase in abundance of senescence cells with increasing chronological age^1,24^. To quantify this in the context of our newly-defined senescence subtypes, we procured a gender-balanced cohort of primary dermal fibroblasts from fifty aging individuals with an age range of 20-89 years old (**Figure 5a**). To confirm the trend of senescence accumulation with age, we first profiled the morphologies of baseline uninduced cells for each of the fifty donors and plotted the relationship between average senescence score with their chronological age. Results showed a significant correlation between the senescence score and the age of the donor with a Pearson correlation coefficient of 0.33 (**Figure 5b**). This trend was conserved for both men and women (**Supplementary Figure 5a-b**) with a Pearson correlation coefficient of 0.32 and 0.34, respectively. With this result, we then asked whether certain senescence subtypes were more strongly associated with chronological age relative to the other subtypes. Plotting the fraction of senescence cells within each senescence subtypes as a function of chronological age, we found that C10 had the strongest age-association with a Pearson correlation coefficient P=0.45, followed by C11 (P=0.30) and C7 (p=-0.16) (**Figure 5c-e**). Furthermore, other morphology clusters also exhibited age-associations, such as C8 (P=0.42) (**Supplementary Figure 5c-l**). Although C10 showed the strongest age-associated relationship, it comprised a minority of the total senescence population, with fractions ranging from 2.5% to 17.5% across all ages tested. C7 on the other hand comprised the majority of the population with a range of 20% to 90%, and C11 ranging from 0.5% to 3.5%. Interestingly, from the initial induction experiments, we found that C7 and C11 were the most dominant senescence subtypes, suggesting that they may be strongly associated with damage responses rather than chronological age. Together, our results indicate that our morphology-based classification of senescence recapitulated the overall age-related accumulation of senescence. However, not all senescence subtypes exhibit the same degree of age-dependence.

**Figure 5.**
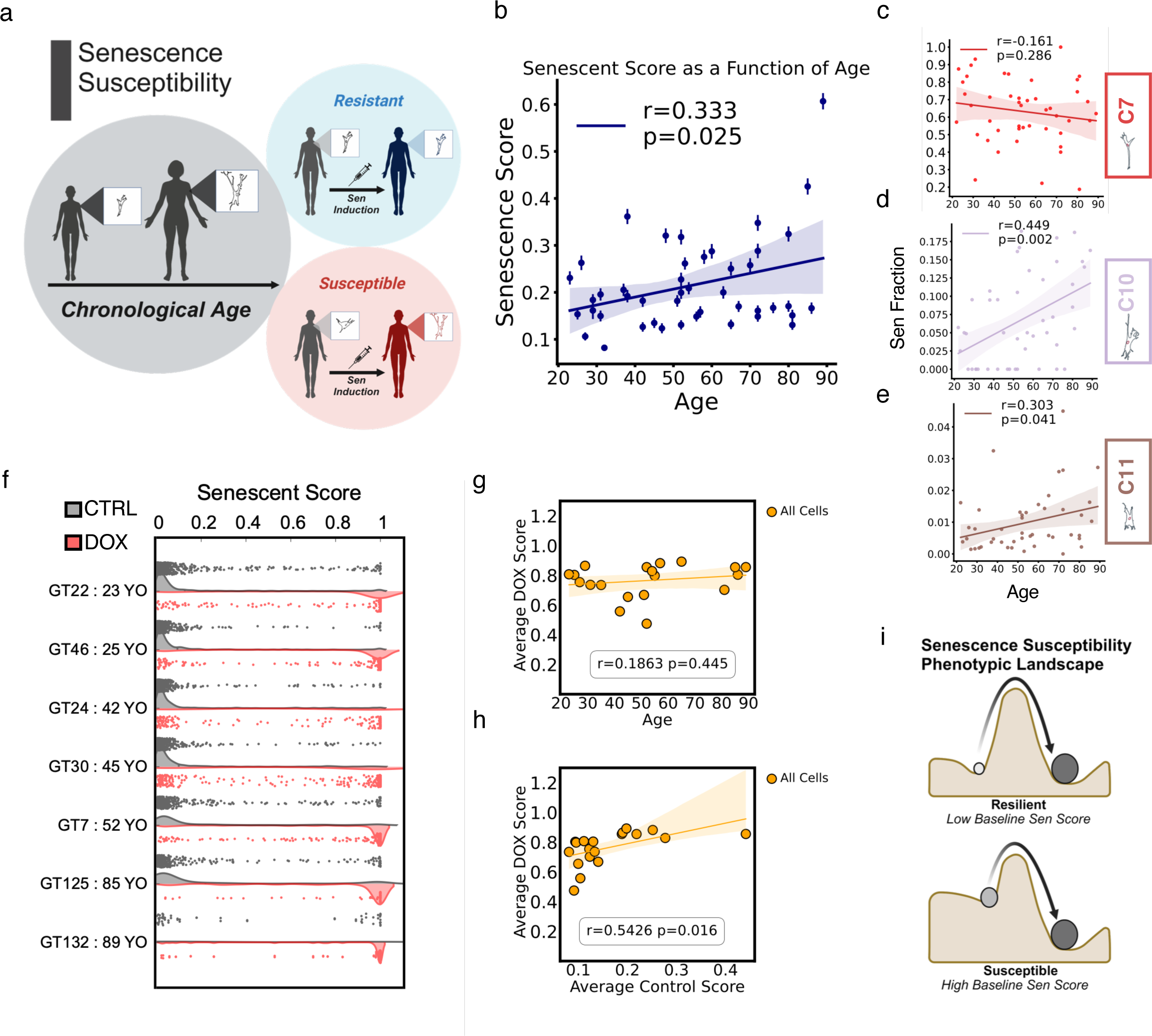
Baseline morphology better encodes senescence susceptibility than chronological age. **a.** Graphical depiction of hypothesized divergent senescent susceptibility responses as a function of age. **b.** Average senescent score from primary dermal fibroblast samples as a function of chronological age fit to a univariate linear regression model (n>300 cells per sample, mean ± SEM). **c-e.** Fraction of senescent cells within each of the three morphological subtypes as a function of chronological age, fit to a univariate linear regression model (from top to bottom: cluster 7,10, and 11 respectively). **f.** Double violin plot for age-dispersed senescent score sample distributions at baseline (grey) and after DOX induction (red) (n>100 cells per condition). **g.** linear regression model of the average DOX-induced senescence score against the chronological age of the sample. **h.** linear regression model of the average DOX-induced senescence score against the average baseline senescent score of the sample. **i.** Senescence susceptibility postulate where induction response is governed by baseline morphological profiles.

Given the relationship between age and senescence score, we wondered whether cells from older donors were more susceptible to senescence induction when exposed to a senescence inducer. To test this, we chose to use DOX, given the clinical relevance to the treatment of cancer. From the same expanded cohort of aging samples, we selected twenty samples that spanned the entire age range from 20-89 and the spread of senescence scores. With each of these aging samples, we induced senescence using the same optimized induction protocols and profiled the senescence score as a function of chronological age (**Figure 5f**, **Supplementary Figure 5m-n**). Plotting the average DOX-induced senescence score versus age, we observed a non-significant correlation, with a Pearson correlation coefficient of 0.19 (**Figure 5g**). Furthermore, looking back at the relationship between the senescence scores and the age, this result is not necessarily surprising given the large donor-to-donor variability in the senescence scores, even for donors of similar chronological ages.

To further elucidate this result, we re-investigated the distributions of senescence scores for cells from each donor, pre and post DOX-induction. Although the spread of distributions recapitulated the variability among donor cells, with a shift towards higher average scores post-induction, we observed an inverse relationship between the spread of the senescence scores and time-point of induction (pre or post). Meaning, that if baseline uninduced cells had a tight but heavy tail distribution of senescence scores, the post-induction scores would have a wider spread of scores (**Figure 5f**. Similarly, if the distribution was wider at baseline, it tended to be tight post-induction. Given this, we considered that if chronological age by itself did not directly associate with senescence scores post induction, whether the baseline senescence score was indicative of senescence score post induction. Plotting the senescence scores at baseline and post induction, we observed a significant correlation with P=0.54, suggesting that the baseline senescence score was indicative of post-induction score, and by extension senescence susceptibility (**Figure 5h**).

Together, our results support the notion that although the senescence burden across cell populations accumulate with increasing age, the relationship between chronological age and senescence susceptibility is more complicated. Given that chronological age positively correlates with baseline uninduced senescence scores, but chronological age is not significantly correlated with senescence score post senescence induction. In any case, the baseline uninduced senescence score is indicative of a cell population’s susceptibility to senescence induction (**Figure 5i**).

### Senescence subtypes exhibit differential responses to Dasatinib + Quercetin

With the emergence of senotherapies as clinically viable strategies to mitigate senescence-associated dysfunctions, we were interested in assessing whether senescence subtypes exhibited differential responses to a small panel of senotherapies. To test this, we profiled the responses of senescent cells to optimized concentrations of five clinically relevant senotherapies comprising both senolytics (D+Q, ARV-825, and Navitoclax (ABT263)) and senomorphics (Fisetin, Metformin) over the course of three days (see Methods). For this experiment, we induced cells from the young donor (GT22) with DOX as previously described, the cultured cells for eight days so that senescence could develop. On day 7, cells were seeded at low density onto Collagen-I coated 96-well glass bottom dishes, and on day 8 cells were stained with live cell markers (SPY650FastAct and SPY555-DNA) to delineate cell and nuclear boundaries) and plates were mounted onto the microscope for live cell imaging. Over the course of 72 hours, images of each condition (5 senotherapies and untreated) were acquired every hour, with 60 hours of post-processing movies used for analysis (see **Methods**). Cells within each of the conditions were segmented at every time point and were assigned to a morphology cluster and senescence subtype after projection into the UMAP space. We chose to perform live cell imaging on the senescent cells exposed to senotherapies specifically to map the dynamic response patterns at single cell resolution over time. Across all experiments we paid particular attention to whether cells died, whether they exhibited plasticity among morphology clusters (*i.e.,* transitioned among senescence subtypes or the other morphology clusters), or whether they were stable in their morphological cluster (**Figure 6a**).

**Figure 6.**
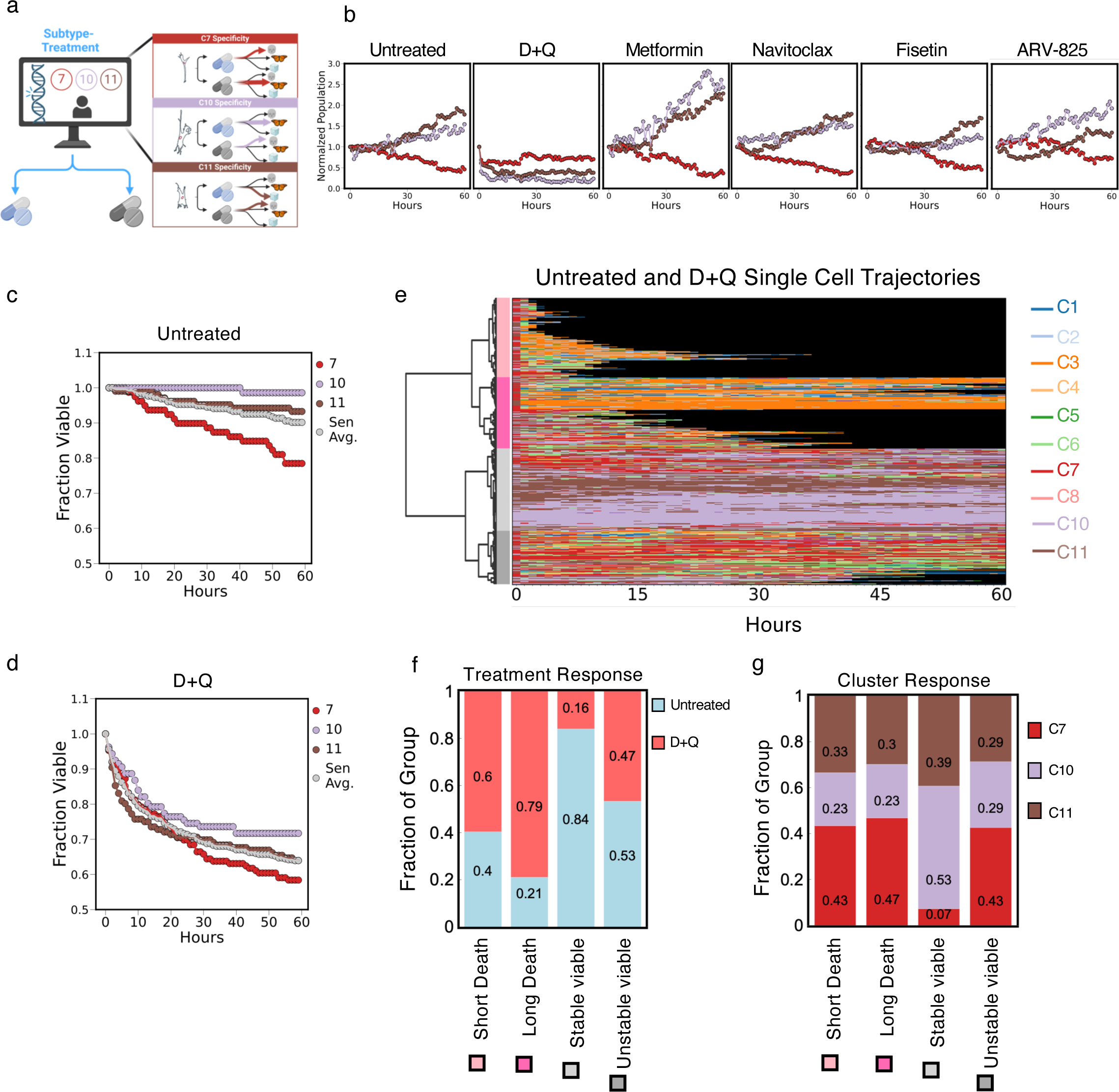
Morphological subtypes encode differential therapeutic response modalities. **a.** Visualization of hypothesized differential therapeutic response profiles of senescence morphological subtypes. **b.** Quasi-population level counts of senescent cell morphological subtypes for untreated, D+Q, metformin, navitoclax, fisetin, and ARV-825 as a function of treatment duration time. Cell number were normalized relative to count at baseline for each condition (t=0 hours, n>150 per subtype per treatment condition)**. c-d.** Single-cell trajectory viability analysis of morphological subtypes as a function of treatment duration time for untreated (c) and D+Q treatment (d). Cells from the induced population were discretized into the different senescence morphological subtypes based on morphology at baseline (t=0, n>100 cells per cluster); cell trajectories were considered non-viable after entering KMEANS clusters 1,2, or 3 and subsequently losing tracking for subsequent frames. Grey *Sen Avg.* line represents the weighted average viability trends. **e.** Heatmap displaying representative morphological evolution of single cells for untreated and D+Q treated starting in one of the three morphological subtypes of senescence. Colors coordinate with KMEANS cluster colors with black indicative of a cell trajectory that has lost tracking (n = 480 cells samples, ward cluster method, Euclidean distance, threshold of 190). Dendrogram clustering was used to determine four overarching response modalities. **f.** Untreated versus Dox enrichment in each of the four response modalities identified in the trajectory heatmap. **g.** Morphological subtype enrichment in each of the four response modalities identified in the trajectory heatmap, normalized to cell count. DOX induced senescent cells in the 23-year old sample were used for this analysis.

With these time-lapse movies, we profiled bulk viabilities across the six tested conditions, which showed a decreased viability for cells treated with D+Q, Navitoclax, and ARV-825, with Fisetin and Metformin having similar viabilities to the untreated controls (**Supplementary Figure 6b**). Additionally, while most of the tested conditions had an average senescence population of ∼60% post induction and treatment, D+Q had a lower fraction of ∼40% (**Supplementary Figure 6c**) and was the only treatment condition showing a bulk decrease in cell count for all three of the senescence subtypes (**Figure 6b**). Given this result and the relevance to ongoing clinical trials of D+Q, we decided to perform deeper analysis on D+Q responses. First, we quantified the fraction of senescence cells at the population level and then within each of the three senescence subtypes as a function of exposure time for the untreated and the D+Q treated cells. Results indicated that there was a decrease of ∼8% of senescence cells over the 60 hours, being largely attributed to spontaneous cell death. C10 and C11 exhibited stable trends over the 60 hours, with C7 exhibiting a decrease to ∼73%, which can be explained by a combination of both spontaneous cell death and transition from C7 to other morphological clusters (**Figure 6c**). On the other hand, D+Q treated senescent cells exhibited ∼45% decrease in the overall fraction of senescent cells, and a complimentary decrease in the abundance of all three senescence subtypes. Interestingly, C10 had a much less decrease after D+Q treatment relative to C11 and C7, with C7 having the largest decrease over the 60-hour treatment (**Figure 6d**).

Quantifying these responses further, we constructed a single-cell response matrix for all senescent cells classified as C7, C10, or C11 at the initial timepoint for both untreated and D+Q conditions. Next, we performed unsupervised hierarchical clustering along the time series to identify response patterns (**Supplementary Figure 6e-f**). Qualitatively, we observed unique and shared response patterns across both conditions. To enable direct comparison among responses for untreated and D+Q senescent cells, we pooled the responses of both conditions then re-clustered the time series data using unsupervised hierarchical clustering. Results yielded four response patterns that were largely defined based on cell viabilities. We named these responses: long death, short death, stable viable, and unstable viable. Short death referred primarily to cells cleared within the first 30 hours, long death were cells cleared after much longer time points (>30 hours), stable viable were cells that remained viable but exhibited little to no transitions to other morphology clusters or other senescence subtypes, and unstable viable were cells that remained viable but exhibited high temporal plasticity and transitions to other morphology clusters or senescence subtypes (**Figure 6e**).

Next, we quantified the abundance of cells within each of the response patterns from the untreated and D+Q conditions. As expected, results showed that senescent cells treated with D+Q made up most of the long and short death classified cells, with 64% and 71%, respectively. Of the cells remaining viable over the 60 hours, the stable viable cells came mainly from the untreated condition (80%, respectively), with 55% of the unstable viable cells coming from the D+Q treatment. These results suggesting that D+Q treatments drove responses of either cell death or instability across the morphological phenotypes (**Figure 6f**). Investigating further the influence of starting senescence subtypes, we observed that C7 was dominate in both the long and short death, comprising 56% and 41% of the cells. C10 was most dominant in the stable viable comprising 54% of cells, and all three of the senescence subtypes were similarly represented in the unstable viable response group (**Figure 6g**). We also found that for cells classified as stable viable starting in C7 exhibited some transitions primarily to C10 and C11, on average spending 28.7% and 27.9% of the time in those clusters, respectively. Cells starting in C10 stayed mainly in C10 (63.4%) and C11 (23.9%), and cells starting in C11 either remaining in C11 (46.2%) or transitioning to c10 (39.9%) (**Supplementary Figure 6i**). Of the unstable viable cells, senescent cells starting in C7 spent most of their time in either C7 (53.7%), C6 (21.2%), or C4 (7.9%). Senescent cells starting in C10 were morphologically very heterogeneous, spending most of the time in either C7 (30.5%), C10 (23.6%), C6 (17%), or C11 (9.3%). Lastly, senescent cells starting in C11 spent most of their time in C6 (37.1%), C11 (20.1%), C7 (19.5%), or C10 (5.1%).

Taken together, results indicate that senescent cells within each of the three senescence subtypes respond differently to D+Q, suggesting that these are indeed functional subtypes of senescence. Notably, senescent cells in C7 were most responsive to D+Q, with cells either dying or becoming morphologically unstable after treatment. C10 senescent cells were most resilient to D+Q treatment. These findings also demonstrate the utility that *ex vivo* morphological profiling and senescence subtyping can be a powerful tool to screen and map response patterns to senotherapies.

## DISCUSSION

Cellular senescence is a heterogeneous phenotype^25^. In summary, we show the development of a robust single-cell approach to classify functional subtypes of senescence (SenSCOUT). Leveraging single-cell profiling with machine learning, we classified a panel heterogeneous aging dermal fibroblast across multiple induction conditions into eleven morphology clusters, out of which we identified three *bona fide* senescence subtypes. Our findings robustly demonstrate that: a) the expressed senescence phenotypes are dependent on the mode of senescence induction and the time post senescence-induction; b) single-cell morphologies encode senescence phenotypes and can be used to determine the protein-based biomarker expressions using imputation via k-nearest neighbor approaches; c) it is not appropriate to assume that all cells exposed to senescence inducers are senescent, since not all cells transform towards senescence and that cell populations can recover (*e.g.,* H2O2) over time post-induction; d) senescence subtypes do not all exhibit similar degrees of age-dependence, with C10 being most strongly associate with chronological age and most resilient to D+Q treatments; e) chronological age by itself does not determine a cell’s susceptibility to senescence induction donor-to-donor variabilities that drives heterogeneity of senescence; and f) senescence subtypes exhibit differential responses to senotherapy D+Q. Together, we provide a robust and promising approach to *ex vivo* senescence profiling and potential applications in precision medicine.

Advances in single-cell technologies and machine learning have enabled deep profiling of cellular phenotypes. Recent studies by other groups applying these techniques to senescence have laid the foundation for identifying senescence and gauging responses to pro-(inducers) and anti-senescence (senotherapies) perturbations^5–7^. However, these studies considered senescence as a binary cellular state (cell as senescence or not) and not as a heterogeneous collection of phenotypes. Building from this, we investigated whether senescence subtypes existed, and whether their existence could help explain the heterogeneous and context-dependence of senescence phenotypes. Importantly, our approach provides deep insights and practical applications of *ex vivo* senescence phenotypes and cellular morphologies. Furthermore, because our appraoch is high-throughput and cost-effective, with the capacity to interrogate both snapshot and dynamic responses, it has direct applications for next-generation screening across a wide range of pro- and anti-senescence perturbations. Furthermore, it can be directly integrated with other image-based or next-generation sequencing-based technologies to identify subtype-dependent molecular vulnerabilities to target senescence phenotypes. Furthermore, coupling with modern spatial multi ‘omic’ technologies can lead to new putative senotherapies and the development of novel approaches, for instance using engineered cell products like CAR-T cells to target senescence subtypes based on specific single cell targets^3,26^.

While our approach hold much promise, one of the current limitations is the fact that it was developed *ex vivo* and have not yet been validated within the context of complex tissue structures. While cells do retain critical memory, for instance of their age and disease status^27^, it will be critical in future studies to investigate the presence and abundance of these senescence subtypes *in vivo* (*i.e.,* within complex skin tissues). To accomplish this, we will need to map senescence subtypes using features that translate well from *ex vivo* to *in vivo*, and *vice versa*. Since the morphologies of cells do not translate directly from cell culture to tissues, we will likely need to use molecular signatures (*i.e.,* transcriptomic, epigenomic, proteomic, etc.) that are established and validated *ex vivo*, then interrogated within tissues (intact or deconstructed) using single-cell or spatial technologies. Furthermore, it will be interesting in future studies to investigate how the presence and abundance of specific senescence subsets alter or contribute to the senescence niche within the aging skin, and how targeting a particular subtype that disrupts the senescence niche may induce beneficial or deleterious effects.

Recent studies have demonstrated the notion of molecular subtypes of senescence^13,28^. In light of these studies and other studies defining molecular signatures of senescence^8,9^, it would be interesting to see what extent these molecular profiles associate with and map onto our morphology-defined subtypes. Furthermore, in our study, we chose to define senescence subtypes as morphology clusters having high average senescence scores (>0.9). While non-senescent clusters (C1, C2, C4, C5) did harbor low senescence scores (<0.3), there were a few clusters, namely C3, C6, and C8 having intermediate senescence scores ranging between 0.6 to 0.82. In future studies it would be interesting to study whether these cells are in fact ‘*poised’* towards senescence, exhibiting a higher tendency towards senescence transformation, and what molecular features best describes these phenotypes. Having a better understanding of this can provide opportunities for targeting precursor phenotypes of which may prove beneficial in certain biological contexts such as aging.

Lastly, another limitation of this approach relates to the potential cell type dependence of senescence phenotypes. Because the senescence subtypes were developed based on the morphologies of dermal fibroblasts, the exact morphologies may only apply to other dermal fibroblasts or other types of fibroblasts/fibroblast-like cells. As with most approaches like this, we will need to investigate the direct utility in other cell types, including fibroblasts from other tissues and organs. While it is almost certainly the case that we will need a new model per cell type (*e.g.,* for epithelial or immune cells), the knowledge learned by establishing the methodology for dermal fibroblast will be directly transferrable. In summary, we present an exciting approach to profile cellular senescence using a robust, cost-effective, and versatile single-cell approach.

## MATERIALS AND METHODS

### Cell culture

Human dermal fibroblasts were procured from the Baltimore Longitudinal Study on Aging (BLSA) and served as the experimental basis for this work. Here we used 50 primary cell samples from a gender-balanced cohort with ages ranging from 23 to 89 years old. Primary cells were cultured under standard conditions on cell culture-treated culture dishes (Corning, 353136) or on Collagen-I coated glass bottom dishes (Cellvis, P24-1.5H-N) with Gibson High-Glucose Dulbecco’s modified Eagle’s medium (DMEM, Thermo Fisher, 11995073) supplemented with 15% fetal bovine serum (FBS, Thermo Fisher, A5256801) and 1% penicillin-streptomycin (Thermo Fisher, 15070063). Cells were grown at 37° Celsius in a humidified incubator with 5% CO2. Cells were passages on average every 3-4 days. Once cultures reached ∼70% confluency, they were plated onto 10 cm cell-culture treated petri dishes (Corning, 430167) at a density of 100,000 cells/dish and allowed to adhere overnight prior to senescence induction. Low passage cells (maximum of 4 passages from stock vial) were used for all experiments except when stated otherwise (*i.e.,* for replicative senescence experiments).

### Induction of senescence

All drugs used for senescence induction were diluted from their respective stock solutes in DMSO (<0.2% vol/vol) according to manufacturer guidelines and subsequently diluted in media to reach the final working concentration for senescence induction (see Table 1 for respective concentrations). Briefly, cell culture media was removed from the petri dishes and replenished with 8 mL of induction media (culture media supplemented with respective concentrations of senescence inducers) and incubated for the specified time durations (see Table 1). Following induction, cells were washed twice with 1X Phosphate-Buffered Saline (PBS, Corning, 21-031) before addition of 8 mL of fresh culture media. The senescent phenotype was allowed to develop, with cellular profiling performed at various time points depending on the experiment; (either 2, 5, 8, 12, or 15 days post induction. For each of the specified induction time points cells were subsequently plated onto 50 ug/mL collagen I (Corning, 354249) coated 24-Well glass bottom plates (Cellvis, P24-1.5H-N) at a density of 1250 cells/well and a volume of 1ml per well to allow for a sparse density and identification of single cell morphologies.

**Table 1.** Senescence induction.

| Method of Induction | Supplier | Concentration | Treatment Duration |
| --- | --- | --- | --- |
| DMSO (VEH Control) | / | 0.5% volume | 4 hours |
| Bleomycin | (SelleckChem, S1214) | 50 μM | 4 hours |
| Hydrogen peroxide | (Millipore Sigma, H1009) | 600 μM | 4 hours |
| Atazanavir | (SelleckChem, S1457) | 80 μM | 96 hours |
| Doxorubicin | (SelleckChem, S1208) | 250 nM | 24 hours |

The end points for replicative senescence per sample (GT22 or GT125) were determined based on a sustained plateau in proliferation rate for a one-week.

### Profiling cellular secretions

To profile the secreted factors from biological conditions of interest, senescent and non-senescent cells were plated at a density of 20,000 cells per 10 cm petri dish and allowed to adhere overnight. The following day, the media was replaced with fresh media and incubated for 48 hours to allow the accumulation of cellular secretions. After 48 hours, the conditioned media was collected, and cells were counted per condition (*i.e.,* per 10 cm dish). Conditioned media was then profiled using Bruker CodePlex Human Adaptive Immune Panel (Bruker Cellular Analysis, CODEPLEX-2L01-4) to quantify the expression of 22 pro-inflammatory markers that overlaps with the documented secretions of senescent cells (SASP—senescence associated secretory phenotype). The raw readouts were calibrated to instrument controls and normalized using the individual cell counts to enable direct comparisons of secreted factors across biological conditions.

### Immunofluorescent staining

Prior to any immunofluorescence staining and quantifications, cells per condition were seeded at low density (1250 cells per well of a 24-well plate) and allowed to completely adhere overnight. Once cells had adhered, they were subsequently washed with 1X PBS and fixed in freshly prepared 4% (w/v) *paraformaldehyde* (PFA, Electron Microscopy Services) for 12 minutes. Cells were then rinsed 3x with PBS before being permeabilized by incubation in a 0.1% Triton-X in PBS solution for 10 min. Cells were again rinsed 3x with PBS to remove any residual Triton-X solution, then blocked with 2% bovine serum albumin (BSA, weight/volume in 1X PBS) for at least 30 minutes at room temperature. Following this blocking step, cells were incubated with various primary antibodies, including p16 (Abcam, ab108349, 1:250), LMNB1 (Abcam, ab229025, 1:500), HMGB1 (Abcam, ab18256, 1:500), or gamma-H2AX (Abcam, ab81299, 1:250) in 1% BSA overnight at 4 degrees Celsius. The next day, cells were washed 3x with PBS and incubated in p21 (Santa Cruz, SC-817, 1:200) in 1% BSA for 1 hour and 15 minutes at room temperature. Cells were washed three times with 1x PBS to remove any residual primary antibodies, then they were subsequently incubated in a cocktail of secondary antibodies and small molecule fluorophores, including Hoechst (Thermo Fisher, H3570 ,1:250), Alexa Fluor-488 phalloidin (Thermo Fisher, A12379,1:200), goat anti-rabbit IgG-Alexa Fluor 568 (Thermo Fisher, A-11036,1:500), and goat anti-mouse IgG-Alexa Fluor 647 (Thermo Fisher, A-21240,1:500) in 1% BSA. Lastly, cells were washed 3x with 1X PBS and imaged using a Leica Stellaris 5 Confocal Microscope. All samples were either imaged immediately after staining or with 24 hours of secondary antibody incubation with intermediate storage at 4 degrees Celsius.

Senescence-associated beta galactosidase fluorescence staining was performed using a similar workflow as previously described. Based on manufacturer’s protocol (Thermo Fisher, C10851) we incubated cells with fluorescence SA β-gal prior to the addition of the p21 primary antibody. Following the incubation with p21 primary antibody, the experimental workflow was consistent with the workflow described above.

### High-content microscopy and quantification of morphological features

Fluorescent images of all conditions were acquired using a Leica Stellaris 5 Confocal Microscope at 20X magnification using 4 laser lines (405 Diode, 488 Diode, 568 Diode, and 647 Diode). Images were taken at 1024 × 1024-pixel resolution with 0.568 microns per pixel. Individual nuclei and cell Boundaries were segmented from raw TIFF files using CellProfiler^TM^ in combination with in-house curation pipelines to ensure well-segmented single cells^17^. Briefly, an immunofluorescence-focused segmentation algorithm used the DAPI stain to delineate the nucleus boundaries and the Phalloidin stain to delineate the general cell shape^29^. Next, we used an in-house curation pipelines to in-silico remove cells from high density regions (more than three toughing cells) to ensure the quantification of single cells. Touching cells made up a very small portion of cells based on optimized cell density and plating. From the generated masks, we computed morphological features primarily describing the sizes, shapes, boundary roughness, and boundary curvatures (**Supplementary Table 1**). We leveraged these masks to also quantify the cellular expressions of protein-based biomarkers of senescence per cell. We profiled >250,000 single cells from snapshot images spanning all the biological conditions presented within this manuscript (not including single cell tracking data).

With the masks generated as outputs from the CellProfiler pipelines we used an in-house feature quantification pipeline to compute ∼218 morphological parameters of each cell^29–31^. These morphological parameters ranged from basic geometric features such as area, perimeter, and curvature to more complex interpretations of morphology such as skeletal features. To identify key parameters driving variance within the cell sample set, we performed primary factor analysis across all cells and all morphological parameters. Parameters expressing a communality value below 0.2 were excluded for subsequent analysis (note a higher communality value indicates a parameter capturing more of the variance across a given cell population) (**Supplementary Figure 2a**). This factor analysis resulted in a reduction of relevant morphological features from 218 to 88 morphological features, which was used for all subsequent analysis.

### Analysis of protein-based biomarkers

Fluorescently labeled protein-based biomarkers were analyzed at single-cell resolution by quantifying the pixel intensity within the cells and nuclei. Specifically, HMGB1 and LMNB1 were quantified using the mean nuclear intensity (integrated intensity per object (*i.e.,* cell or nuclei) divided by the total number of pixels (*i.e.,* object size), p21 was quantified by total nuclear expression (*i.e.,* summation of the pixel intensities per nuclei), and p16 and βGAL were quantified by total cellular expression (*i.e.,* summation of the pixel intensities per cell, included nuclear region). To correct of any effects from potential batch -to-batch variations we used ReCombat^TM^ software for all datasets across biological replicates. Protein biomarker expressions for all cells across induction condition were log-transformed prior to z-score normalization. The control distributions for each cell were used to compute the Z-Score values for cells based on their individual protein expression (performed per age sample).

To determine the relative cutoffs for senescence (senescent vs. non-senescent cells), we plotted the distributions of cells for baseline (mostly non-senescent cells) and induced cells, then identified the intersection point that defined the cross-over for each of the distributions, *i.e.,* cells with low values in the baseline distributions but were high in the induced conditions were denoted as a senescence threshold per biomarker. This enabled us to determine the average intersection point of the control distribution and the distribution of H_2_O_2_ and Bleomycin senescence induction conditions for the five biomarkers (Bleomycin and H_2_O_2_ were selected as they were strong, orthogonal inducers). We determined these values to be approximately 1.6 for young (GT22) and 1.3 for old (GT125), with the older having a smaller z-score cutoff likely due to more baseline cells expressing the senescent phenotype at baseline. Note these values correlate for approximately 10% of the healthy GT22 cells being classified as senescent and 15% for GT125 cells.

### UMAP and k-means clustering

We constructed the reduced dimensionality framework using Uniform manifold approximation and projection (UMAP) across all biological conditions and replicates (n=3) for GT22 (23-year-old) and GT125 (89-year-old). All morphological parameters were independently log-normalized and standard scaled (z-score). This “preprocessed” morphological parameter dataset was subsequently used to construct a 2D-UMAP space. UMAP is a nonlinear dimensionality reduction algorithm that capturer and projects the structure of high dimensional data in a lower-dimensionality space (for this work, the 88-vector space was reduced to two with Euclidean convergence). Each point in the UMAP space represents an individual cell. Given UMAP is a nonlinear reduction across multiple morphological parameters, there is no ‘equivalent’ with each of the constituent vectors, however UMAP-1 correlated well with cell and nuclear size and UMAP-2 with cell and nuclear shapes. As a complement to the UMAP morphological analysis, we performed k-Means clustering to discretize and identify distinct morphologically-defined groups within the dataset. We performed k-means clustering analysis using eleven cluster, which was determined based on the inertia and silhouette values. Here the goal was to identify the number of clusters that would minimize the inertia (typically at the elbow) and maximize the silhouette values (*i.e.,* stability of the respective clusters). Across the eleven clusters we identified *bona-fide* senescent and non-senescent cell populations, however, cluster 9 harbored a large fraction of mis-segmented cells. For initial analysis cluster 9 was kept for cell classification but was dropped from analysis when reporting actual findings of the data. To quantify morphological heterogeneity, the Shannon entropy was calculated using the following equation for the ten validated k-means clusters (*i.e.,* eleven clusters with the removal of cluster 9).

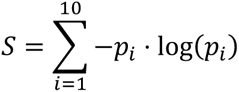

Here *S* is the Shannon entropy (greater magnitude signifies a more heterogeneous population) and *p_i_* is the fraction of cells classified within morphological cluster *i* (here *i* range from 1-10).

### Protein biomarker imputation

Given the interconnectedness of morphology and senescence-associated protein biomarker expression, we developed and implemented a computational pipeline to determine biomarker expression based on cell and nuclear morphologies parameters alone. This is particularly useful since using our imaging workflow, we can stain for a maximum of 2 biomarkers at a time. This biomarker imputation workflow enabled us to determine the expression of 5 biomarkers per single cell across all the biological conditions of interest. Principal component analysis was performed on the 88 log-normalized, standard-scaled morphological parameters that was used to construct the 2D-UMAP and identify the 5 principal components capturing 95% of the sample variance. For a given cell with an unknown biomarker, the 20 nearest neighbor cells that were stained for that biomarker if interest were identified from the weighted Euclidean distance of the 5 principal components. For computational streamlining, nearest neighbors were restricted to the same cell line (GT22 or GT125) within the same k-means clusters. After nearest neighbors were identified, we computed the expression using a weighted average of the biomarker. The weights were determined by Euclidean distance to the imputed cell, with cells in closer proximity being weighted more. These steps were iterated for each of the biomarkers.

### Kinetics of senescence experiments

Human dermal fibroblast samples were induced with either H_2_O_2_ or doxorubicin, and the senescence phenotypes were allowed to develop over 2, 5, 8, 12, or 15 days in separate dishes. At these time points, cells were seeded at low density on collagen-I coated 24-well glass-bottom plates. Cells were fixed, immunofluorescence stained for senescent associated protein biomarkers, and imaged as described above. This data was analyzed for biomarker expression behavior and morphological cluster enrichment as a function of senescence induction time.

### Machine learning approach to quantify senescence subtypes using Xception algorithm

XCEPTION, a 48-layer convolutional neural network architecture for image classification was used to identify non-parameterized trends in morphological signatures associated with senescence. This architecture was largely based on the original namesake, with two key supplements and 1,000,000 total trainable parameters: (1) weights were initialized from ImageNet and (2) additional pooling, dropout, and SoftMax layers were appended to the end of the architecture (to allow for a senescence probability as a final output of the model). To construct the input samples to the model, a 3-layer dataset consisting of 1536 x 1536-pixel images were created for individual cells where the first layer was the textured nuclei image (grayscale H33342 image), the second layer was the textured actin architecture (grayscale Phalloidin image), and the third layer was the composite overlay of both the nuclei and actin (combined H33342 and Phalloidin images). Each image was geometrically centered and had pixel intensities normalized between 0-1. This dataset was constructed from the same cells used to create the UMAP manifold (GT22 and GT125 dermal fibroblast samples across the different induction conditions). Every layer of the input sample set was overlaid against a neutral background (pixel value of zero). Each sample of the training dataset was tagged as either senescent or non-senescent based on protein biomarker expression z-score cutoffs (described above). Specifically, cells differentially expressed in both biomarkers were tagged as senescent whereas cells not differentially expressed in either biomarker were tagged as healthy.

Given the surplus of senescent-tagged cells compared to controls (because of 5 senescent induction conditions as opposed to one control), the training set was subsampled from the larger dataset at large to balance both senescent and healthy cells. Furthermore, the training set was balanced for age and the senescent-tagged cells were further balanced for induction conditions to minimize potential bias. Non-senescent tagged cells were biased towards control samples (>80% of samples were control conditions in either age group). A generative model was applied to augment the dataset by implementing non-distortive transformations (i.e. flips and rotations) to maintain the integrity of size and shape effects of the training set. Training of the model was conducted in two parts, with an 80-20 train-test split. The first 200 epochs were trained with ImageNet initialized weights, frozen backbone, Adam’s Learning rate of 1×10^−4^, and accuracy-based optimization. Next, the backbone was unfrozen and trained for an additional 20 epochs with an Adam’s Learning Rate of 1×10^−3^ which allowed for a finer convergence of algorithm accuracy. The accuracy of model along with the area under the curve (AUC) of the receiver operating curve (ROC) were used to evaluate model performance.

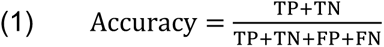

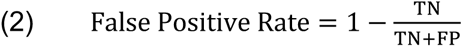

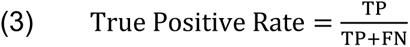

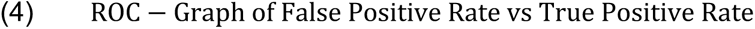

Here, TP is true positive, TN is true negative, FP is false positive, and FN is false negative.

### Evaluating the effects of aging on senescence experiments

Forty-eight additional primary dermal fibroblast samples of known gender and chronological age were cultured and fixed at low densities using experimental workflows described above. Morphologies of each patient at baseline were analyzed and we determined their senescent score and their enrichment within morphologically-defined senescence subtypes. Using a univariate linear regression model we quantified how the abundance of senescence cells and the senescence score correlated with donor age. 18 samples were further selected to assess chronological age-associated response to senescence induction. Samples were either treated to DMSO or Doxorubicin (8-day induction period) and subsequently fixed for morphological analysis. Morphologies of cells in both conditions were analyzed by the trained Xception model for score distributions and analyzed by the k-means algorithm to determine senescent subtype distribution. Using this two-pronged orthogonal approach, we quantified how the abundance of senescence within each of the three bona-fide senescence subtypes changes with increasing donor age.

### Senotherapy response and live-cell imaging

GT22, human dermal fibroblast samples were induced using a 8-day doxorubicin induction protocol and plated at a density of 300 cells/well in a 96 well Collagen-I coated glass-bottom plate (Cellvis, P96-1.5H-N). After allowing the cells to adhere overnight, wells were replenished with fresh media containing Spirochrome 555-DNA (Spirochrome, SC201,1:1000) and Spirochrome 650-FastAct (Spirochrome, SC505 ,1:1000) and incubated for four hours at physiological conditions in a humidified cell culture incubator. A baseline series of image was taken of the cells prior to treatment with serotherapeutic drugs. Following the acquisition of the baseline images, wells were supplemented with either Dasatinib (SelleckChem, S1021, 1 µM) and Quercetin (SelleckChem, S2391, 10 µM), Metformin (SelleckChem, S1950, 5 µM), Navitoclax (SelleckChem, S1001, 5 µM), Fisetin (SelleckChem, S2298, 25 µM), or ARV-825 (SelleckChem, S8297, 10 nM) as a one time, bolus treatment while plates were on the microscope. Images were taken once every hour for a span of 60 hours to capture the response dynamics per drug. Similar to the fixed images, cells in each frame were segmented using an optimized CellProfiler^TM^ workflow, and the trackpy Python package was used to ascribe cell identities across frames of the same imaged well. Segmented images were analyzed using the same UMAP and k-means analysis described above to determine the morphological clusters of individual cells as a function of time.

### Dasatinib +quercetin single-cell responses and viability

Dasatinib+Quercetin treatment was used as the focus for single-cell viability as it was the most effective senotherapy based on cell killing from our small senotherapy screen. To determined dead or dying cells based on the time-lapse movies, two criteria were assessed based on its morphological progression: (1) the cell has lost tracking and (2) in the frame before the cell has lost tracking, it must be in either cluster 1, 2, or 3 (indicative of a senescent dying phenotype). Although clusters 1, 2, and 3 were identified as non-senescent clusters at baseline, cells belonging to these clusters after senescence induction and senotherapy treatment exhibited enhanced expression of DNA dama as denoted by gamma H2AX. Cells were classified by their initial morphological subtype using the pre-treatment images and we analyzed their as a function of senotherapy time to assess subtype-specific responses. Given an uneven number of cells in each senescent morphological cluster, we normalized cell counts for each subtype. To identify how cell morphology transitions as function of treatment, a hierarchal clustering algorithm was fed a dataset of all senescent morphological subtype trajectories (*i.e.,* a cell with a baseline starting in one of the senescent morphological subtypes (C7, C10, C11) for both untreated and D+Q treatment conditions). The hierarchical clustering used the ward algorithm to discretize four response patterns: short death, long death, stable viable, and unstable viable.

### Statistics and reproducibility

To establish statistical comparisons between multiple groups we performed One-way ANOVA and post-hoc pairwise Tukey tests. Correlations of linear regression fits were evaluated using R and p-values, respectively. Data was log-transformed, where applicable, to normalize distributions for gaussian-based models/ No data was excluded from this work. The experiments were not randomized, and the authors were not blinded to the study. All experiments were performed with at least 3 biological replicates with at least 2 in-plate technical replicates.

### Software

All single cell segmentations were performed using optimized workflows in CellProfiler. All dynamic tracking data was performed using a custom algorithm combining CellProfiler outputs with TrackPy algorithum. All analyses were performed in Python with the following software specifications: python 3.8.12, tensorflow-gpu 2.5.0, scipy 1.7.1, scikit-learn 1.1.1, pandas 1.4.3, matplotlib 3.4.2, seaborn 0.12.0, plotly 5.10.0, trackpy 0.5.0, umap-learn 0.5.3, and numpy 1.21.2.

## Code availability

The source code used to generate the data presented in this work was deposited on the Phillip Lab GitHub page

## Data availability

All data supporting the findings of this study are available within the paper its Supplementary Information, or the Phillip lab GitHub page. All other data (including large image files and movies) are available from authors upon reasonable request.

## Acknowledgements

We acknowledge the financial support for this work from American Federation for Aging Research (AFAR)— Glenn Foundation Junior Faculty Award (JMP), The Johns Hopkins University Older Americans Independence Center of the National Institute on Aging under award number P30AG021334 (JW, JMP), The National Institutes of Health 1UG3CA275681-01 (PW, JMP), Start-up funds from the Biomedical Engineering Department and the Whiting School of Engineering at Johns Hopkins University (JMP), and National Institute of General Medical Sciences of the National Institutes of Health under Award Number R35-GM142889 (JF)

## Author contributions

P.K, J.W, and J.M.P conceived and designed study; P.K, N.M, Y.L, A.W, L.P, T.S, B.S performed experiments; P.K, N.M, J.F, C.M, J.M.P conceived and performed formal analysis; P.K, J.F, P-H.W developed analysis algorithms and software; P.K, N.M, J.F, J.M.P interpreted results; P.K, N.M, J.M.P wrote and edited manuscript; All authors edited and contributed to the final manuscript

## Conflict of interest

P.K, N.M., J.M.P are inventors on a patent application related to this work. All other authors declare no conflict of interest.

## Notes

### Summary of Updates

I revised the order of the author list to have me appear last.

