## Supplementary Figures for "Single-cell morphology encodes functional subtypes of senescence in aging human dermal fibroblasts" (from another version of this preprint: version 1, 2024-05-14)

# S1

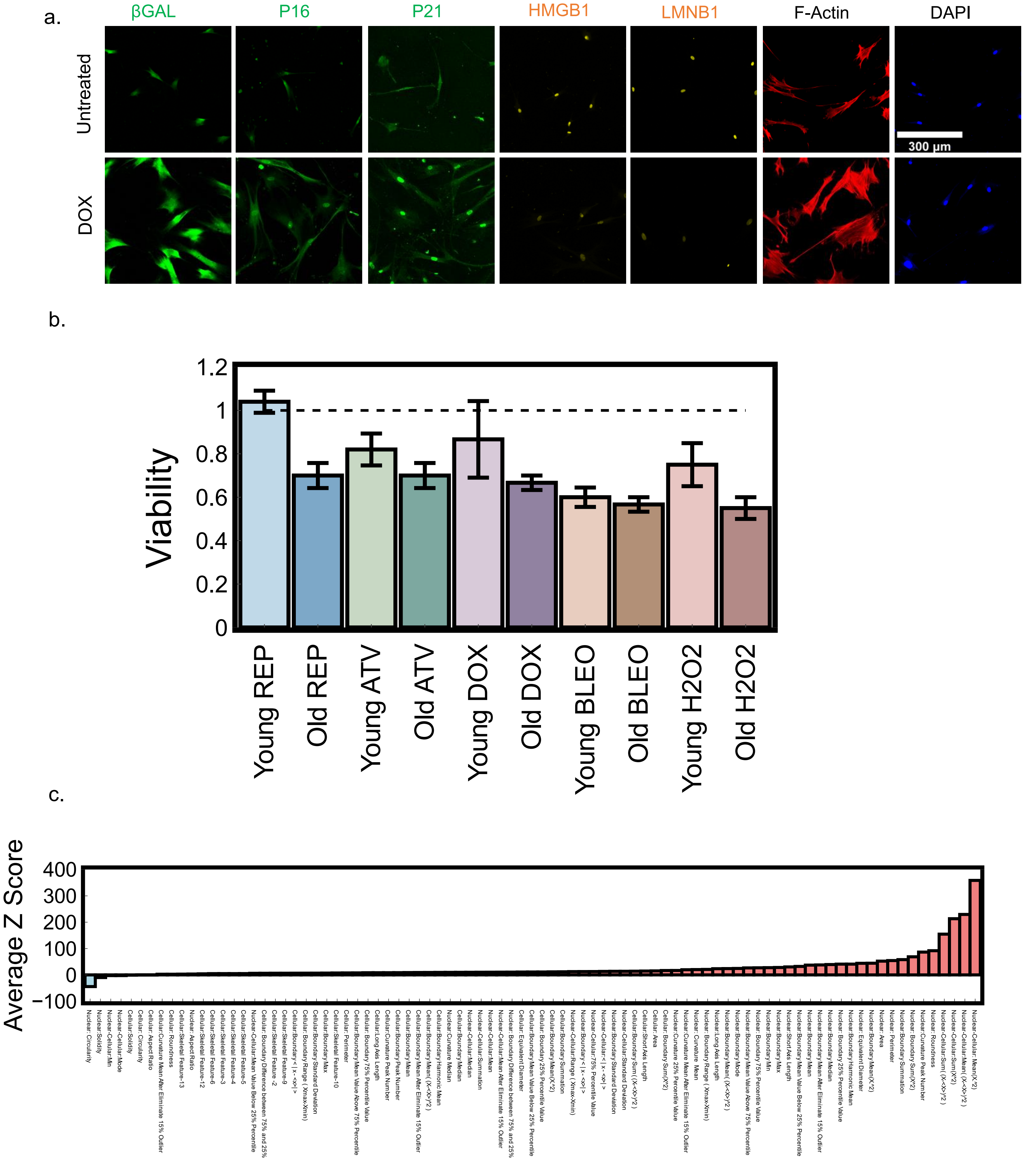
a.
βGAL
P16
P21
HMGB1
LMNB1
F-Actin
DAPI
Untreated
DOX
300 µm
b.
Viability
1.2
1
0.8
0.6
0.4
0.2
0
Young REP
Old REP
Young ATV
Old ATV
Young DOX
Old DOX
Young BLEO
Old BLEO
Young H2O2
Old H2O2
c.
Average Z Score
400
300
200
100
0
−100

### S1-Caption

**Fig. S1 | a** Representative images of the immunofluorescent staining of 5 senescence-associated protein biomarkers across untreated (DMSO vehicle) and DOX-induced primary dermal fibroblast samples (age 85-Male). **b** Viability of senescence induction conditions 8 days post induction. Values less than one indicate cell death, values greater than one indicate cell proliferation (n=3 biological repeats, Mean ± SEM). **c** Waterfall plot of all cellular and nuclear morphological features enrichment to either untreated or senescence-induced samples across both ages.

S2

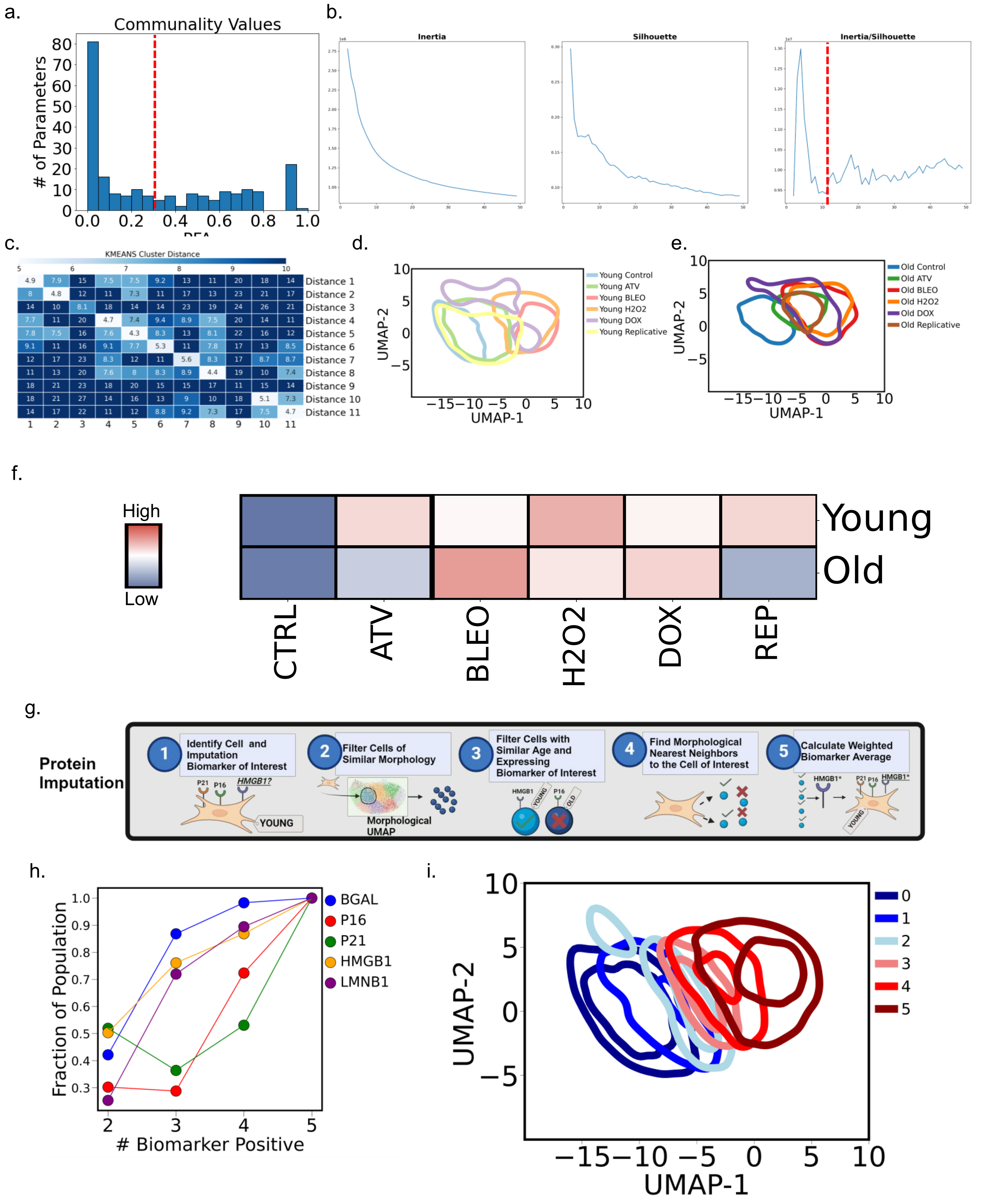
a.
Communality Values
### of Parameters
80
70
60
50
40
30
20
10
0
0.0
0.2
0.4
0.6
0.8
1.0
b.
Inertia
Silhouette
Inertia/Silhouette
c.
KMEANS Cluster Distance
5
6
7
8
9
10
4.9 7.9 15 7.5 7.5 9.2 13 11 20 18 14 Distance 1
8 4.8 12 11 7.3 11 17 13 23 21 17 Distance 2
14 10 8.1 18 14 14 23 19 24 26 21 Distance 3
7.7 11 20 4.7 7.4 9.4 8.9 7.5 20 14 11 Distance 4
7.8 7.5 16 7.6 4.3 8.3 13 8.1 22 16 12 Distance 5
9.1 11 16 9.1 7.7 5.3 11 7.8 17 13 8.5 Distance 6
12 17 23 8.3 12 11 5.6 8.3 17 8.7 8.7 Distance 7
11 13 20 7.6 8 8.3 8.9 4.4 19 10 7.4 Distance 8
18 21 23 18 20 15 15 17 11 15 14 Distance 9
18 21 27 14 16 13 9 10 18 5.1 7.3 Distance 10
14 17 22 11 12 8.8 9.2 7.3 17 7.5 4.7 Distance 11
1 2 3 4 5 6 7 8 9 10 11
d.
UMAP-2
UMAP-1
Young Control
Young ATV
Young BLEO
Young H2O2
Young DOX
Young Replicative
e.
UMAP-2
UMAP-1
Old Control
Old ATV
Old BLEO
Old H2O2
Old DOX
Old Replicative
f.
High
Low
Young
Old
CTRL
ATV
BLEO
H2O2
DOX
REP
g.
Protein Imputation
1 Identify Cell and Imputation Biomarker of Interest
P21
P16
HMGB1?
YOUNG
2 Filter Cells of Similar Morphology
Morphological UMAP
3 Filter Cells with Similar Age and Expressing Biomarker of Interest
HMGB1
YOUNG
P16
OLD
4 Find Morphological Nearest Neighbors to the Cell of Interest
5 Calculate Weighted Biomarker Average
HMGB1*
P21
P16
HMGB1*
YOUNG
h.
Fraction of Population
1.0
0.9
0.8
0.7
0.6
0.5
0.4
0.3
2
3
4
5
### Biomarker Positive
BGAL
P16
P21
HMGB1
LMNB1
i.
UMAP-2
UMAP-1
0
1
2
3
4
5

### S2-Caption

**Fig. S2 | a** Primary factor analysis communality histogram distribution, approximate cutoff at 0.2. **b** Inertia, silhouette, and inertia/silhouette graphs to determine optimal number of k-means clusters. Optimal number selected from plateau in inertia/silhouette without introducing too many clusters, resulting in 10 total. **c** Heatmap of cluster stability by average distance to centroid for cells within each cluster. Cluster 9, the mis-segmented cluster, had a dispersed cluster proximity, indicating cluster instability. **d,e** Single contour topology overlaid on UMAP space for experimental conditions for young (**d**) and old (**e**) samples (One contour per condition). **f** Heatmap of Shannon entropies for each condition from the morphological k-means clusters. Higher Shannon entropy values correlate with higher heterogeneity. 'High' Denotes the maximum Shannon entropy value of 2.3 while low denotes a value of 1.7. **g** Experimental workflow for the protein imputation platform. **h** Fraction of population positive for a specific biomarker given all biomarkers determined through imputation (positive determined by z-score difference from control of 1.58 for young and 1.3 for old, respectively). For example, if a cell has 3 biomarkers meet their respective senescence cutoffs, βGAL has a 90% probability of being one of those biomarkers whereas p21 has a 30% chance. **i** Contour map demonstrating morphologic location of cells depending on number of biomarkers positive (positive determined by z-score difference from control of 1.58 for young and 1.3 for old, respectively). The ground truth value was known for two biomarkers with the remaining three determined through the imputation platform.

S3

a.

| | Young BLEO Z SCORE | Young H2O2 Z SCORE | Old BLEO Z SCORE | Old H2O2 Z SCORE |
|---|---|---|---|---|
| P16 | 1.173291893 | 1.262087093 | 0.926382744 | 0.96835085 |
| HMGB1 | 1.272657019 | 1.285656265 | 1.107763853 | 1.149055524 |
| LMNB1 | 1.263464231 | 1.145878251 | 1.134212362 | 1.173647252 |
| BGAL | 1.199683123 | 1.465753156 | 1.042658545 | 1.14685942 |
| P21 | 1.555928099 | 1.202053631 | 1.00416376 | 0.890842252 |
| Mean | 1.293004873 | 1.272285679 | 1.043036253 | 1.06575106 |

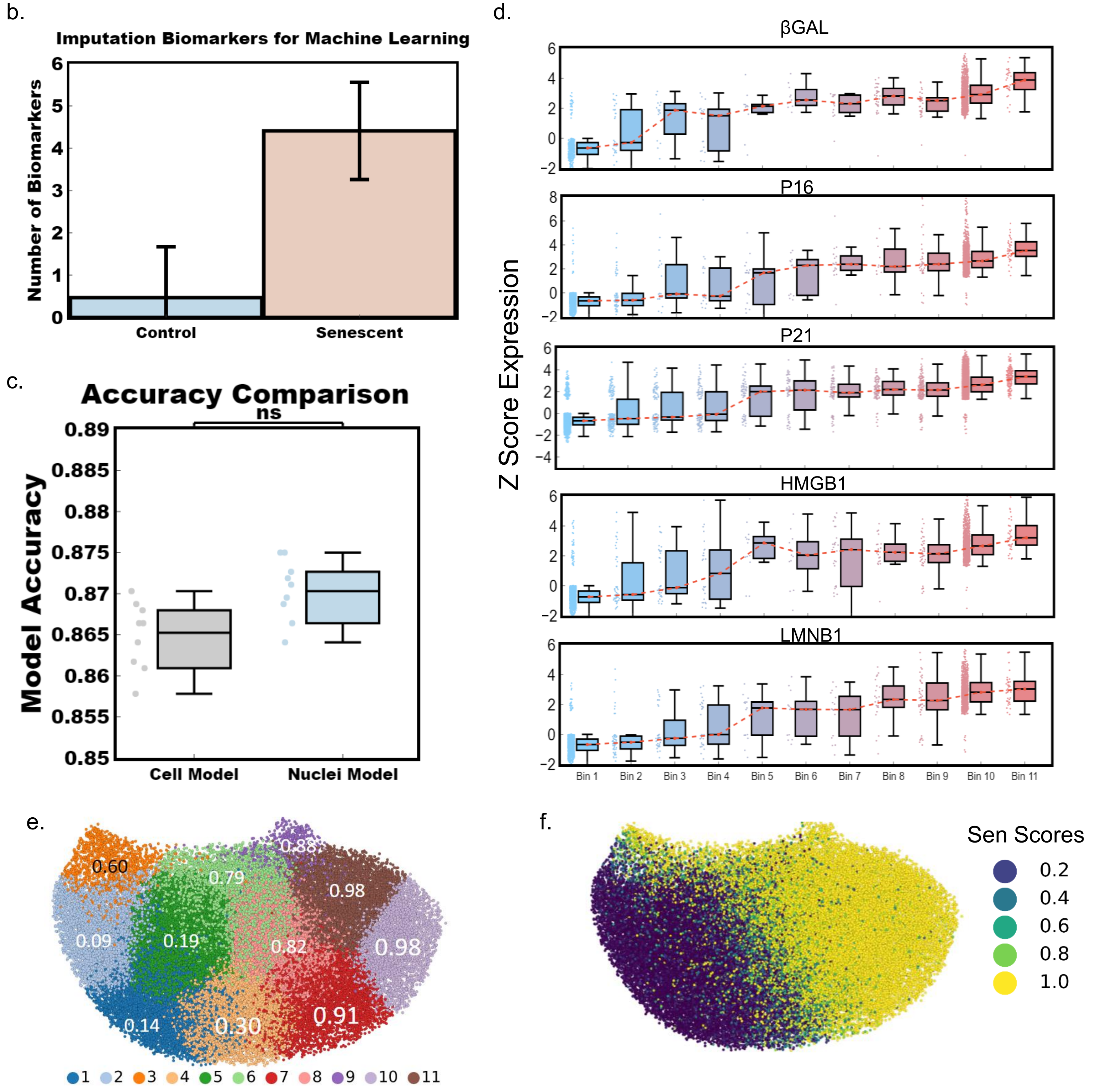

### S3-Caption

**Fig. S3 | a** Log-normalized distribution intersection between control and BLEO/H2O2 induction conditions for both young and old samples across all biomarkers.  The average intersection was then padded by a 5% buffer to create senescent cutoffs for each age. **b** Average amount of biomarkers positive in ‘non senescent’ (control) and ‘senescent’ classification fed into the Xception machine learning architecture ( n>500, mean ± STDEV). Dataset was constructed only on the two ground truth biomarkers stained for each cell; this graph imputes the remaining three to construct the distribution. **c** Accuracy of nuclei- and cell- only models on the same dataset as the composite model (with a mean accuracy of 89%). **d** distribution of Z-score expression of each biomarker in 0.04 interval bins ranging senescent scores from 0-1 (mean ± 95% CI). **e**  Mean senescence score of each k-means cluster overlayed on top of the UMAP space. **f** Individual senescent scores overlayed on top of the UMAP space.

S4

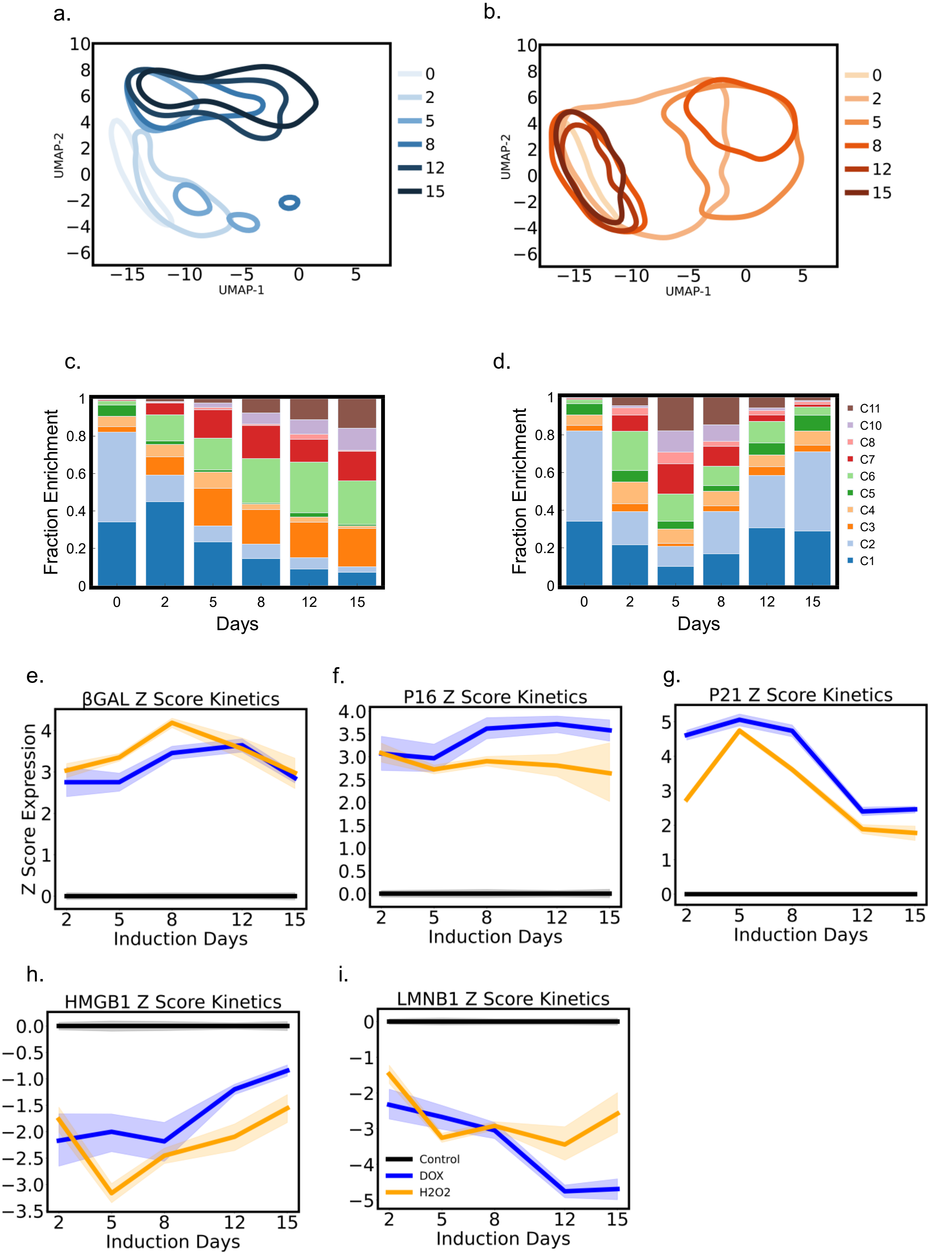

a.
b.
UMAP-2
UMAP-1
0
2
5
8
12
15
c.
d.
Fraction Enrichment
Days
C11
C10
C8
C7
C6
C5
C4
C3
C2
C1
e.
βGAL Z Score Kinetics
f.
P16 Z Score Kinetics
g.
P21 Z Score Kinetics
Z Score Expression
Induction Days
h.
HMGB1 Z Score Kinetics
i.
LMNB1 Z Score Kinetics
Control
DOX
H2O2

### S4-Caption

**Fig. S4 | a,b** Morphological contour distributions as a function of induction time for DOX- (**a**) and H2O2- (**b**) induced cells. **c,d** Stacked bar plot for k-means cluster enrichment as a function of induction time for DOX- (**c**) and H2O2- (**d**) induced cells. **e-i** Average Z-score quantification of βGal (**e**), p16 (**f**), p21 (**g**), HMGB1 (**h**), and LMNB1 (**i**) as a function of time post senescence induction (shading indicates 95% CI around the mean). This analysis only used cells within one of the three morphological subtypes (k-means clusters 7,10, and 11). 23-male fibroblasts were used for this analysis.

# S5

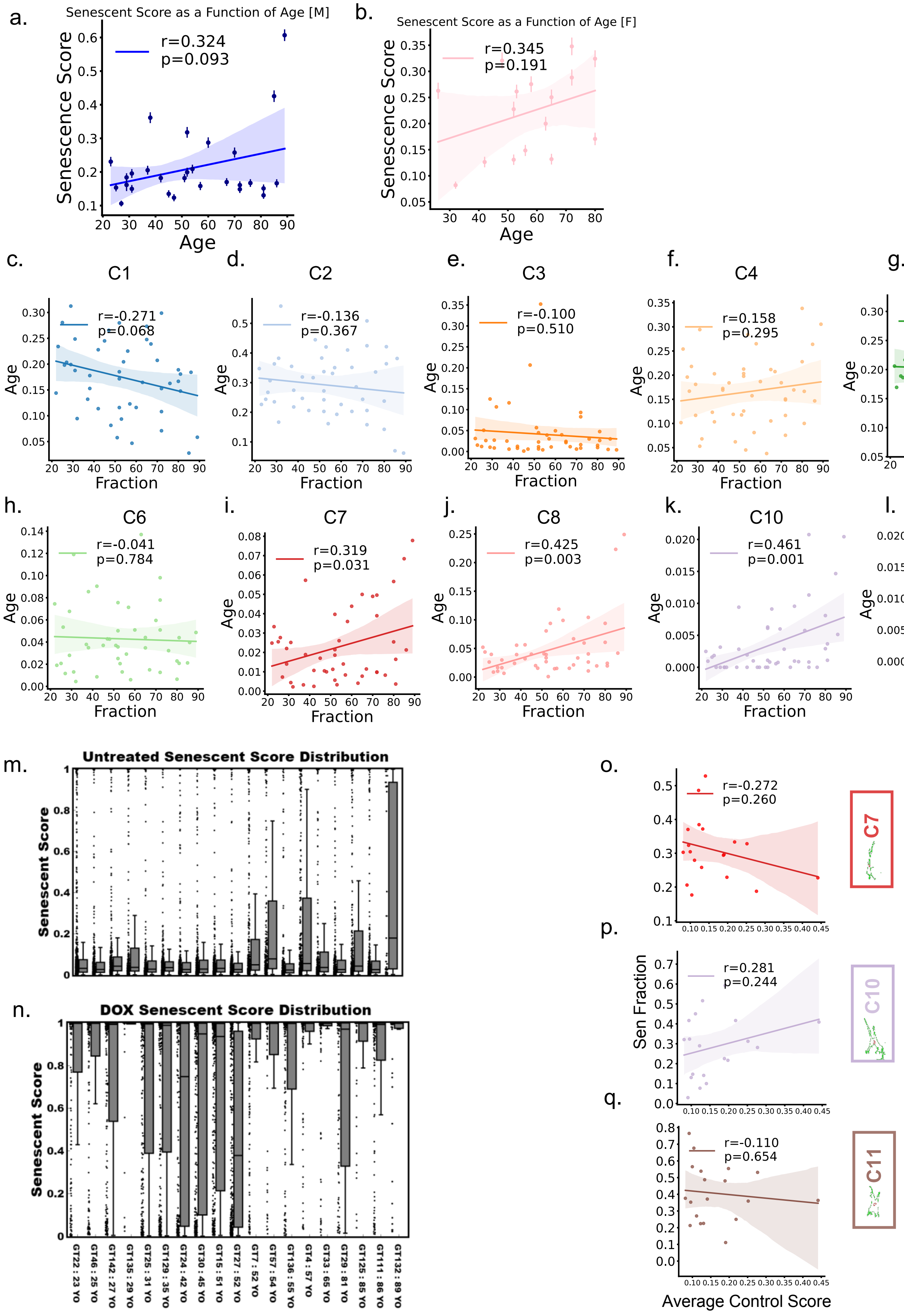
a.
Senescent Score as a Function of Age [M]
r=0.324
p=0.093
Senescence Score
Age
b.
Senescent Score as a Function of Age [F]
r=0.345
p=0.191
Senescence Score
Age
c.
C1
r=-0.271
p=0.068
d.
C2
r=-0.136
p=0.367
e.
C3
r=-0.100
p=0.510
f.
C4
r=0.158
p=0.295
g.
C5
r=-0.041
p=0.785
h.
C6
r=-0.041
p=0.784
i.
C7
r=0.319
p=0.031
j.
C8
r=0.425
p=0.003
k.
C10
r=0.461
p=0.001
l.
C11
r=0.303
p=0.041
Age
Fraction
m.
Untreated Senescent Score Distribution
Senescent Score
n.
DOX Senescent Score Distribution
Senescent Score
GT22 : 23 YO
GT46 : 25 YO
GT142 : 27 YO
GT135 : 29 YO
GT25 : 31 YO
GT129 : 35 YO
GT24 : 42 YO
GT30 : 45 YO
GT15 : 51 YO
GT27 : 52 YO
GT7 : 52 YO
GT57 : 54 YO
GT136 : 55 YO
GT4 : 57 YO
GT33 : 65 YO
GT29 : 81 YO
GT125 : 85 YO
GT111 : 86 YO
GT132 : 89 YO
o.
r=-0.272
p=0.260
C7
p.
r=0.281
p=0.244
C10
Sen Fraction
q.
r=-0.110
p=0.654
C11
Average Control Score

### S5-Caption

**Fig. S5 | a,b** Average Xception senescent score from primary dermal fibroblast samples as a function of chronological age fit to a univariate linear regression model for male (**a**) and female (**b**) (n>300 cells per sample, mean ± SEM). **c-l** Fraction of cells within each of the k-means clusters as a function of chronological age, fit to a univariate linear regression model (numerically ascending cluster numbers from left to right). **m** Box and whisker plot of the baseline (untreated) senescent score distribution of 18 selected samples. **n** Box and whisker plot of the DOX-induced senescent score distribution of 18 selected samples. **o-q** Fraction of senescent cells within each of the three morphological subtypes post DOX-induction as a function of the average baseline senescent score, fit to a univariate linear regression model (from top to bottom: cluster 7,10, and 11 respectively).

S6-1

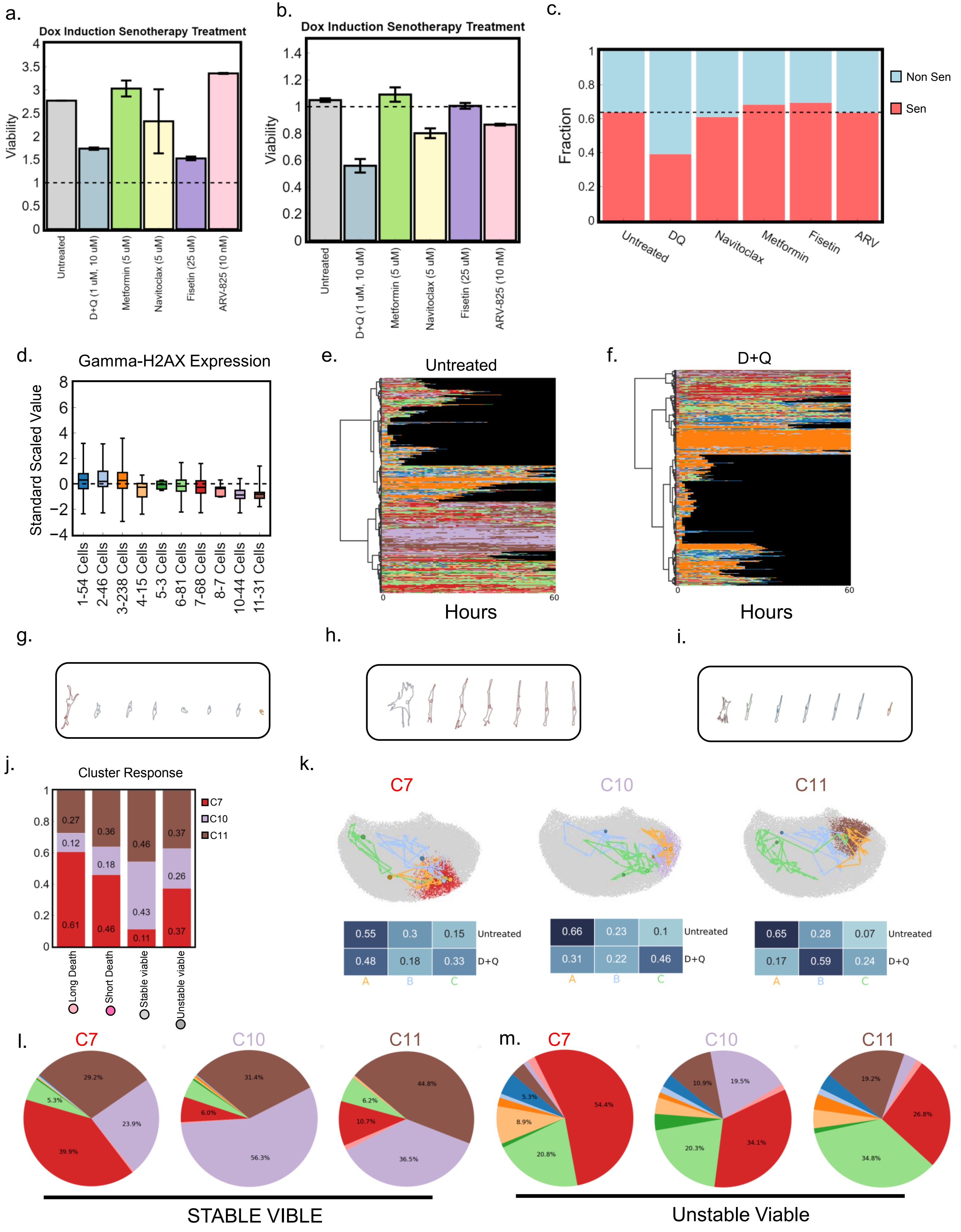

a.
Dox Induction Senotherapy Treatment
Viability
Untreated
D+Q (1 uM, 10 uM)
Metformin (5 uM)
Navitoclax (5 uM)
Fisetin (25 uM)
ARV-825 (10 nM)
b.
Dox Induction Senotherapy Treatment
Viability
c.
Fraction
Non Sen
Sen
Untreated
DQ
Navitoclax
Metformin
Fisetin
ARV
d.
Gamma-H2AX Expression
Standard Scaled Value
1-54 Cells
2-46 Cells
3-238 Cells
4-15 Cells
5-3 Cells
6-81 Cells
7-68 Cells
8-7 Cells
10-44 Cells
11-31 Cells
e.
Untreated
Hours
f.
D+Q
Hours
g.
h.
i.
j.
Cluster Response
C7
C10
C11
Long Death
Short Death
Stable viable
Unstable viable
k.
C7
C10
C11
Untreated
D+Q
A
B
C
l.
m.
STABLE VIBLE
Unstable Viable

### S6-Caption

**Fig. S6 | a,b** Viability of control (**a**) and DOX-induced (**b**, 8-day induction) to a three-day treatment to various senotherapeutic agents. Values less than one indicate cell death, values greater than one indicate cell proliferation (n=3 biological repeats, Mean ± SEM). **c** Fraction of cells in senotherapeutic treated DOX-induced cells within one of the three morphological subtypes (7,10,11- 'Sen') or not ('Non sen'). **d** Standard scaled mean nuclear gamma-H2AX expression for morphological KMEANS clusters. Dataset was comprised of both untreated and D+Q treated, DOX-induced cells. **e,f** Heatmap displaying representative morphological evolution of single cells for untreated (**e**) and D+Q treated (**f**) for any starting cluster (n>300 single cells per treatment , ward cluster method, Euclidean distance) . Colors coordinate with KMEANS cluster colors with black indicative of a cell that has lost tracking. **g-i** senescent morphological subtype representative morphological responses to D+Q for subtype 7 (**g**), 10 (**h**), and 11 (**i**); 5-hour intervals taken per cell starting from time zero. j morphological subtype enrichment in each of the four response modalities identified in the trajectory heatmap, not normalized. **k** Three representative time-series k-means morphological transitions trained from untreated and D+Q conditions (top, subtypes 7,10, 11 from left to right). Abundance of each time series k-means patterns for untreated and D+Q treatment treatments for each morphological subtype (bottom). **l** time spent in each morphological k-means cluster for cells originating in one of the senescent subtypes, in the *stable viable* response group (originating in subtype 7,10,11 left to right). **m** time spent in each morphological k-means cluster for cells originating in one of the senescent subtypes, in the *unstable viable* response group (originating in subtype 7,10,11 left to right). DOX induced senescent cells in the 23-male sample were used for this analysis.

S7

a.

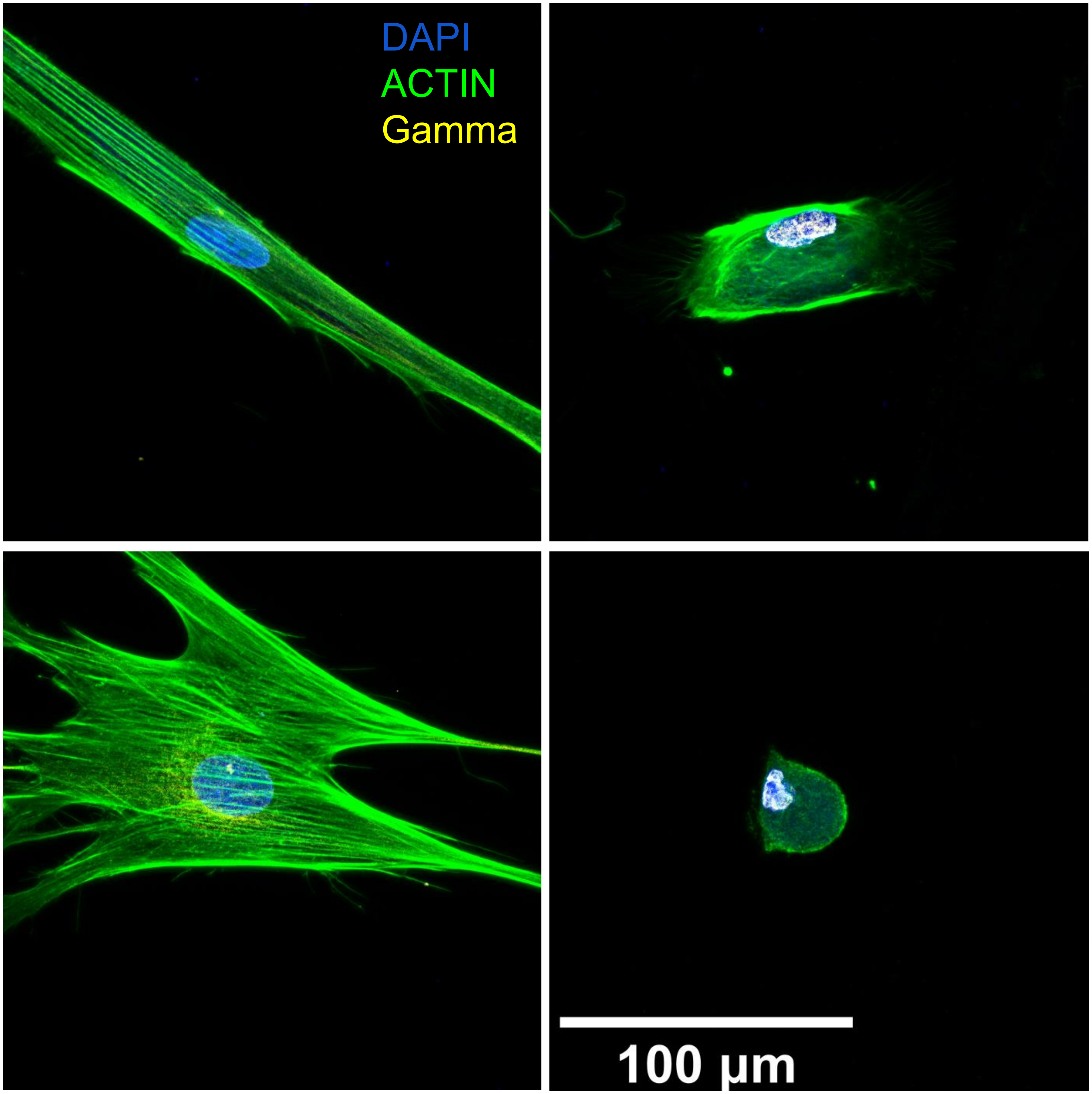

### S7-Caption

**Fig. S7 | a** 60X representative images of Gamma-H2AX Foci in DOX-induced cells (23-male) to D+Q treatment. Unperturbed senescent morphologies (left) express less damage foci than smaller morphologies (right).
